# Orthogonal control of gene expression in plants using synthetic promoters and CRISPR-based transcription factors

**DOI:** 10.1101/2021.11.16.468903

**Authors:** Shaunak Kar, Yogendra Bordiya, Nestor Rodriguez, Junghyun Kim, Elizabeth C. Gardner, Jimmy Gollihar, Sibum Sung, Andrew D. Ellington

## Abstract

**Background:** The construction and application of synthetic genetic circuits is frequently improved if gene expression can be orthogonally controlled, relative to the host. In plants, orthogonality can be achieved via the use of CRISPR-based transcription factors that are programmed to act on natural or synthetic promoters. The construction of complex gene circuits can require multiple, orthogonal regulatory interactions, and this in turn requires that the full programmability of CRISPR elements be adapted to non-natural and non-standard promoters that have few constraints on their design. Therefore, we have developed synthetic promoter elements in which regions upstream of the minimal 35S CaMV promoter are designed from scratch to interact via programmed gRNAs with dCas9 fusions that allow activation of gene expression.

**Results:** A panel of three, mutually orthogonal promoters that can be acted on by artificial gRNAs bound by CRISPR regulators were designed. Guide RNA expression targeting these promoters was in turn controlled by either Pol III (U6) or ethylene-inducible Pol II promoters, implementing for the first time a fully artificial Orthogonal Control System (OCS). Following demonstration of the complete orthogonality of the designs, the OCS was tied to cellular metabolism by putting gRNA expression under the control of an endogenous plant signaling molecule, ethylene. The ability to form complex circuitry was demonstrated via the ethylene-driven, ratiometric expression of fluorescent proteins in single plants.

**Conclusions:** The design of synthetic promoters is highly generalizable to large tracts of sequence space, allowing Orthogonal Control Systems of increasing complexity to potentially be generated at will. The ability to tie in several different basal features of plant molecular biology (Pol II and Pol III promoters, ethylene regulation) to the OCS demonstrates multiple opportunities for engineering at the system level. Moreover, given the fungibility of the core 35S CaMV promoter elements, the derived synthetic promoters can potentially be utilized across a variety of plant species.

## Introduction

The field of synthetic biology aims to revolutionize biotechnology by rationally engineering living organisms (1–6). One aspect of rational engineering is to embed biological organisms with complex information processing systems that can be used to control phenotypes (2, 3, 7, 8), often via synthetic gene circuits that can predictability regulate and tune expression of endogenous as well as transgenes (4, 9–11).

However the performance of such synthetic genetic circuits is often plagued by unwanted interactions between the circuit components and the host regulatory system, which can lead to loss of circuit function (10). These unprogrammed interactions can be mitigated via the design and use of genetic parts that have minimal cross-talk with the host, creating orthogonal regulatory or orthogonal control systems (OCS) that can further serve as the basis for constructing complex genetic programs with predictable behaviors. In the last two decades an increasing number of well-characterized genetic parts have been combined in circuits capable of complex dynamic behaviors, including bi-stable switches, oscillators, pulse generators, Boolean-complete logic gates (7, 12–15). While OCS and the circuits that comprise them were initially characterized in microbial hosts, more recently a significant fraction of them have been constructed and characterized in eukaryotic hosts such as yeast and mammalian cells (12, 16–19). More recently, synthetic transcriptional control elements have begun to be characterized in plants (20–22).

While a variety of artificial plant transcription factors containing diverse DNA binding domains and plant-specific regulatory sequences are known (20, 22), orthogonal control requires more programmable DNA binding domains and modular regulatory domains (20, 22–24). To this end, we describe an alternate strategy for the construction of orthogonal transcriptional regulatory elements in plants, powered by a single universal transcriptional factor – dCas9:VP64 which has been shown to work in a wide variety of eukaryotic species, including plants (16, 25, 26). While this transcription factor has primarily been used for the regulation of endogenous genes (25–27), here we describe a generalizable strategy for the universal design and use of synthetic promoters that rely only on the production of specific gRNAs to program dCas9:VP64, and the use of this set of mutually orthogonal promoters for the bottom-up construction of circuits that show multiplexed control of gene expression.

### Design of a modular cloning framework for facile construct assembly

Traditionally the process of construction of these synthetic gene expression systems has relied on time-consuming practices of recombinant DNA technology like design of custom primers, PCR amplification, gel extraction of PCR products. Over the last decade the advent of high-throughput cloning techniques, such as Golden-gate cloning with Type IIS restriction enzymes, has greatly accelerated the design-build-test cycle for the construction and prototyping of synthetic circuits (7, 9, 28, 29). Because the overlaps for assemblies can be modularly specified, multiple parts can be assembled sequentially in a single tube reaction.

While a Golden-Gate framework was previously described for the construction of plant expression vectors (30), here we used the highly optimized modular cloning (MoClo) framework, instantiated as a yeast toolkit (YTK) as the basis of our architecture (28). Recently, beyond yeast expression vectors, this framework has been adapted for the construction of a mammalian toolkit (MTK) (9). Along with both YTK and MTK, a plant toolkit based on the YTK architecture will prove essential for seamlessly porting parts and circuits across diverse eukaryotes. Briefly, in this framework the entire vector is divided into particular ‘part’ types flanked by BsaI restriction sites followed by a unique ligation site. Promoters, genes and terminators are generally categorized into Type 2, 3 and 4 parts respectively where each part type has a unique overhang that dictates the compatibility between part types (9, 28) (**Fig 1A, S1A**). This preserves the architecture of each transcriptional unit (promoter-gene-terminator). For the assembly of multiple transcriptional units (TU), each transcriptional unit is first cloned into an ‘intermediate’ vector flanked by connector sequences that dictate the order of the TUs to be stitched together. By using appropriate connectors, each TU can be further assembled into a final expression vector in a single pot reaction (**Fig S1B**) [20]. This modular approach enables rapid assembly of increasingly complex genetic circuits comprised of multiple transcriptional units.

**Figure 1.**
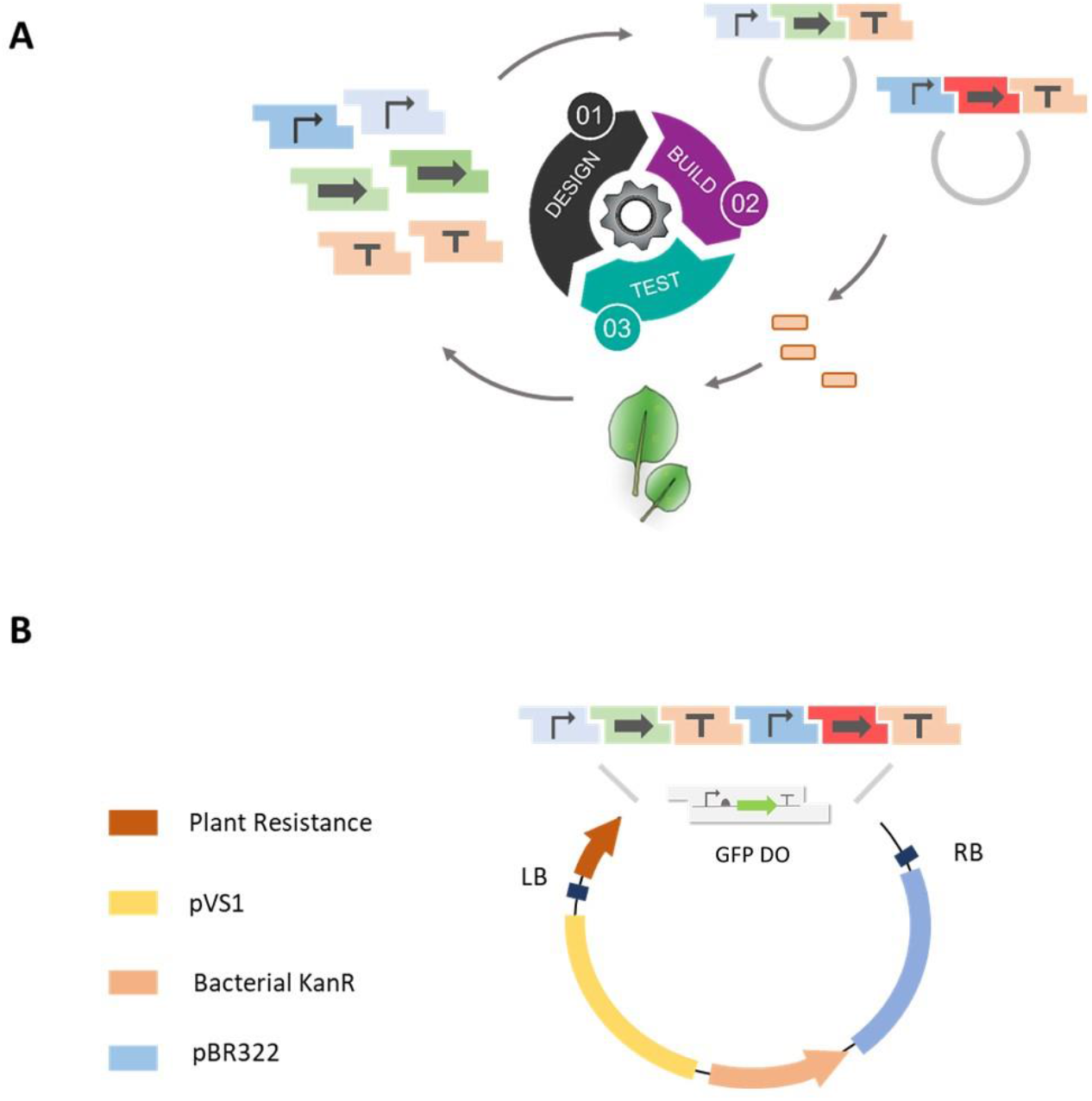
Schematic overview of the design-build-test cycle. **A**. Genetic elements such as promoters, genes and terminators are encoded as modular parts consisting of BsaI recognition sites flanked by specific overhangs to ensure the hierarchical assembly of transcriptional units. Once assembled, the constructs are transformed into Agrobacterium and the reporter expression is characterized in *Nicotiana benthamiana* leaf infiltrates **B**. Design of the shuttle vector backbone used for the assembly of constructs and subsequent propagation in *Agrobacterium*.

Since *Agrobacterium*-based transformation has been the staple for plant genetic engineering for decades (31), we used compatible vectors as the basis for our framework, and designed and constructed three YTK-compatible shuttle vectors. Each expression vector contains the pVS1 replicon (an Agrobacterium origin of replication – OriV and two supporting proteins – RepA and StaA) and pBR22 origin for propagation in *Agrobacterium* and *E.coli* respectively, and a common antibiotic selection cassette (KanR) that has been shown to be functional in both species (**Fig 1B, Materials and Methods**) (29, 30). The three constructs otherwise differed in the plant selection marker - BASTA, hygromycin, and kanamycin. The resistance markers were expressed from the Nos promoter and also contained a Nos terminator (30) (**Fig 1B**). The backbone also contains a GFP drop-out cassette that allows easy identification of correct assemblies, which should appear as colonies that lack fluorescence (9, 28) (**Fig 1B**).

Fluorescence and luminescence reporters are frequently used to study protein localization and interaction in plants and animals (32). To provide these useful reporter parts in the context of our system, we cloned the strong promoter from Cauliflower mosaic virus (35S) as a Type 2 part and its corresponding terminator as a Type 4 part (33, 34). These parts can be matched with a number of fluorescent reporter genes (GFP, BFP, YFP and RFP) all as Type 3 parts for robust reporter expression. Combinations of these proteins can also potentially be used for BIFC (Bimolecular Fluorescence Complementation) (35). Similarly, luciferase is commonly used in plant molecular biology to study circadian rhythm (36), test the spatiotemporal activities of regulatory elements (37), and to study the plant immune system (38, 39). Therefore we adapted a luciferase gene from *Photinus pyralis*, commonly known as firefly luciferase (F-luc) (21).

Single TUs comprised of a 35S promoter, fluorescent reporter genes and the luciferase gene, and a terminator that serves as a polyadenylation signal were assembled into the *Agrobacterium* shuttle expression vector (**Fig 2A-C**). The activity of constructs was assayed using transient expression in *Nicotiana benthamiana* (30). As expected, we see strong activity of the promoter with the expression of the respective reporter genes (**Fig 2A-C**). In order to diversify the promoters used in circuits (and thereby avoid recombination and potentially silencing), we also included a well-characterized promoter from the Ti plasmid that drives mannopine synthase (Pmas) (40–43). When the 35S promoter was swapped with Pmas, similar expression levels of YFP were achieved (**Fig 2D**).

**Figure 2.**
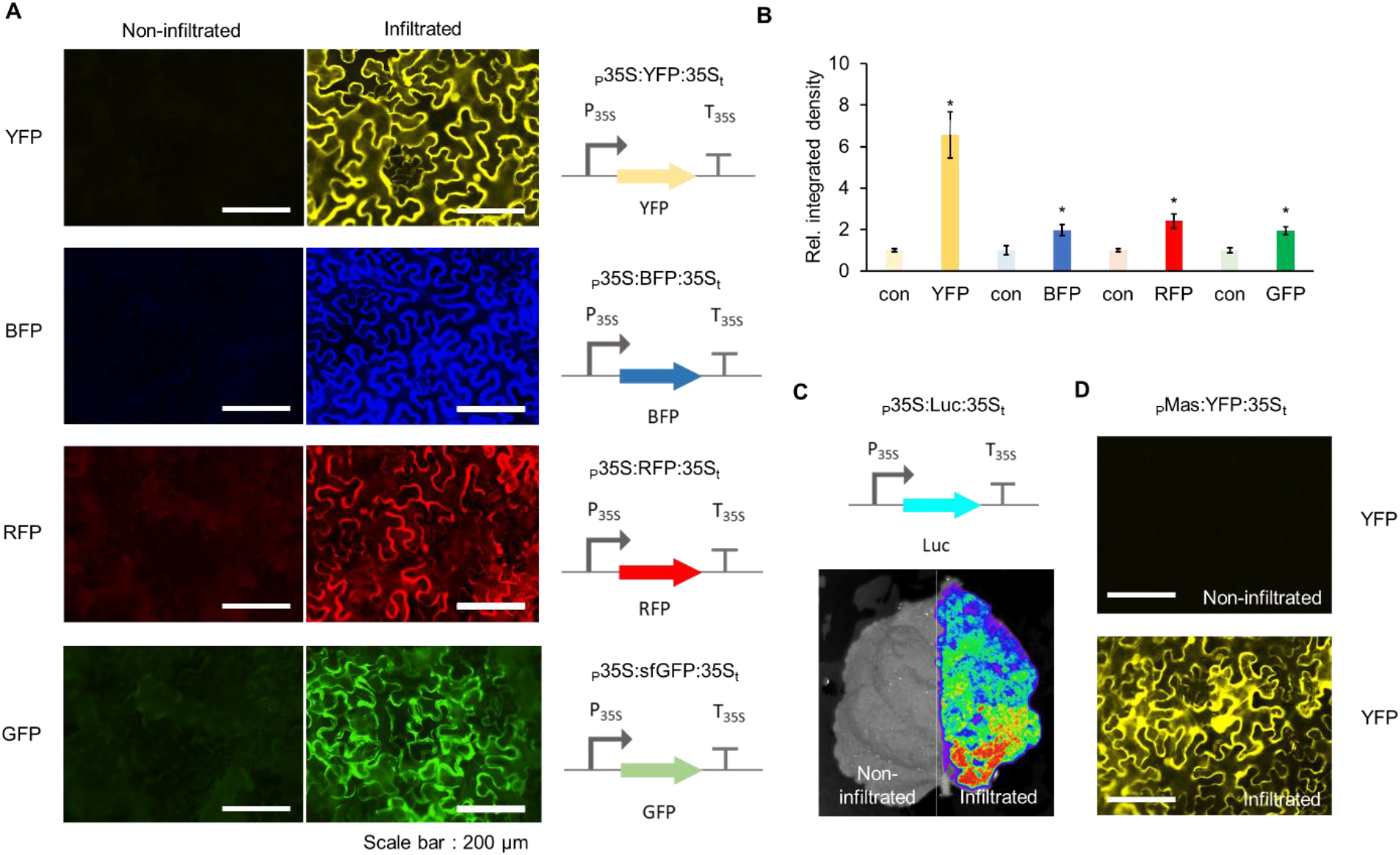
Characterization of reporter constructs assembled using APT toolkit. **A**. Fluorescence microscope images showing *Agrobacterium* mediated transient expression of YFP, BFP, RFP and GFP under the control of 35S promoter into *Nicotiana benthamiana* leaves. Images on the left are from non-infiltrated leaves (negative control) captured using the appropriate filter at same exposure and gain settings as was used for the positive images on the right (**Material and Methods**). **B**. Relative integrated density of each fluorescence signal (shown in panel A). Integrated density was measured using image J software and normalized to that of a non-infiltrated control (con). Error bars: S.D. (n=3, independent replicates). Asterisks indicate statistical significance in a student t-test (P<0.05). **C**. Luminescence reporter luciferase expression shown by *Agrobacterium* mediated transient expression of luciferase in *Nicotiana benthamiana* leaves. Left half of the leaf was not infiltrated with *Agrobacterium*. **D**. Fluorescence microscope images showing *Agrobacterium* mediated transient expression of YFP under MAS promoter in *Nicotiana benthamiana* leaves. Image on the left is the brightfield image for the same construct.

### Development of an Orthogonal Control System (OCS) to regulate transgene expression

One of the primary difficulties with using synthetic biology principles and methods to engineer organisms, especially in eukaryotes, is that the functionality of synthetic circuits is often plagued by unwanted interactions of the circuit ‘parts’ with the underlying regulatory machinery of the host (44). As a particularly relevant example, systems developed in the past for transgene expression caused severe growth and developmental defects in *Arabidopsis* and *Nicotiana benthamiana* (45, 46). Therefore, it is paramount to develop regulatory tools to control transgene expression that minimizes the impact on endogenous plant machinery/physiology, while maintaining the modularity and scalability of synthetic approaches in general.

A potential solution to this problem is to develop orthogonal ‘parts’ that of necessity function independently of endogenous regulation by the host. To this end, we set out to develop a fully integrated Orthogonal Control System (OCS) based on orthogonal synthetic promoters driven by an Artificial Transcription Factor (ATF). We started with the deactivated form of the Cas9 protein (dCas9) fused to the transcriptional activator domain VP64 as a highly programmable ATF (26, 27). While dCas9:VP64 has previously been shown to upregulate the expression of endogenous genes via specific guide RNAs (gRNAs) that target the promoter region upstream of those genes (25, 47), this strategy has not been utilized for the construction of a fully orthogonal system in which custom promoters can be similarly regulated. Here we develop a suite of synthetic promoters (pATFs, promoter for Artificial Transcription Factor) in which each promoter has a similar modular architecture: varying number of repeats of gRNA binding sites followed by a minimal 35S promoter (33, 34). This system is inherently scalable, since new binding sites bound by new gRNAs can be built at will. The complete list of parts (promoters, genes and terminators) is provided in **Supplementary Table 1**.

We initially varied the number of gRNA binding sites (3 and 4) upstream of the minimal 35S promoter, and analyzed expression of the reporter using transient assay in *Nicotiana benthamiana*. Three repeats provided the best expression of the reporter gene without significant background (**Fig 3A**). The promoter architecture was further assayed for leaky expression by generating pATF:YFP/BFP/RFP constructs and expressing gRNA constitutively in the absence of dCas9:VP64 (**Fig 3A**). None of these constructs show expression above background (**Fig 3B and 3C**). However, upon the addition of constitutively expressed dCas9:VP64 cassette to the circuit, induction of reporter protein expression was observed (**Fig. 3B and 3C**). Each pATF demonstrated comparable levels of expression (pATF1:YFP - 3-fold, pATF3:BFP - 6-fold and pATF4:RFP - 2 fold) compared to that obtained from the regular 35S promoter (6-fold; **Fig 2B**). The basic features of the pATF and corresponding gRNAs can thus form the basis for the OCS and should allow us to predictably control reporter and other gene circuits. The complete list of assembled OCS circuits is provided in **Supplementary Table 2**; as the reader will see, OCS circuitry can be organized in terms of increasing complexity and demonstrates how the Design-Built-Test approach can be used to empirically generate ever more substantive plant phenotypes.

**Figure 3.**
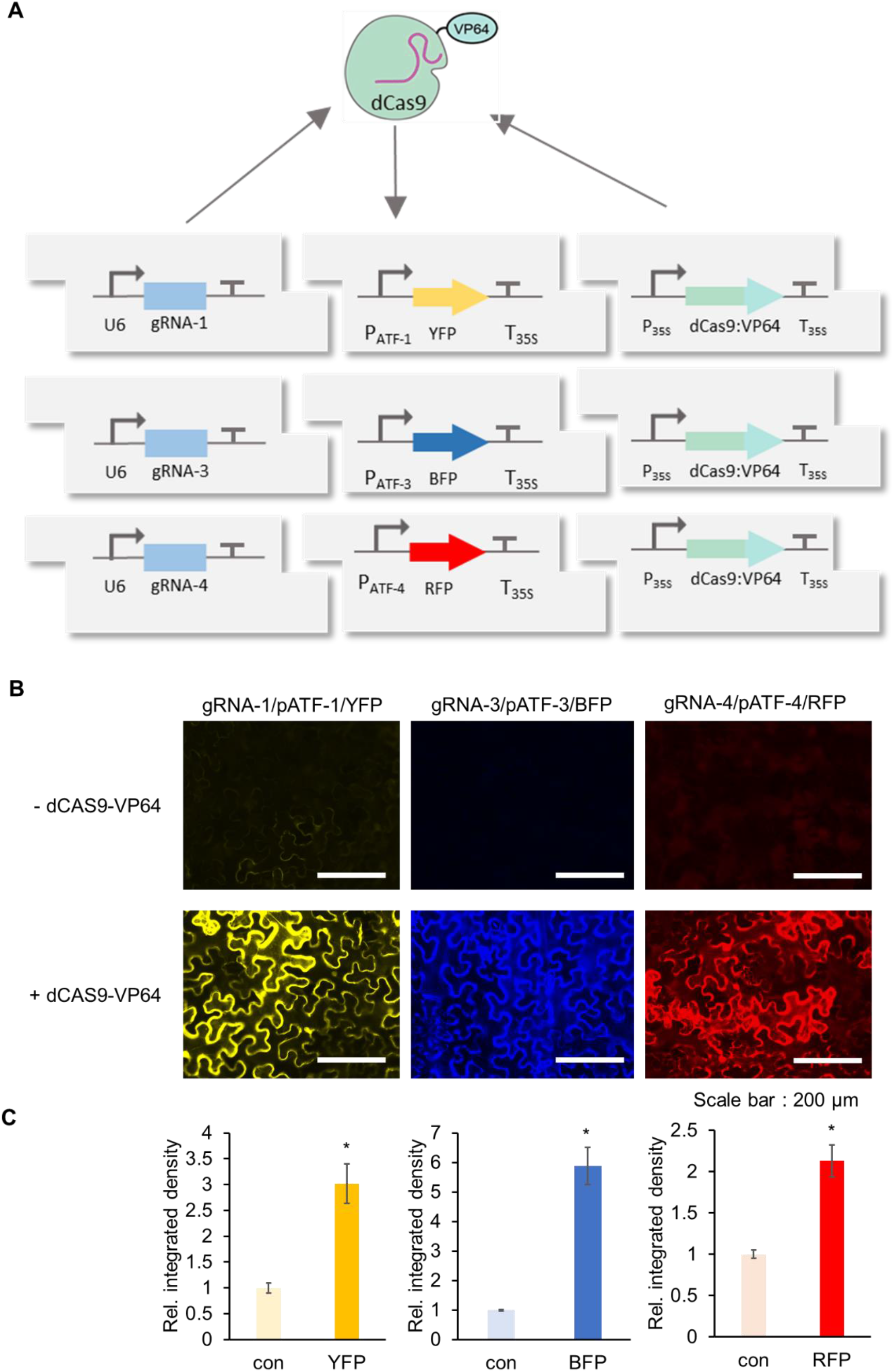
Characterization of activity of synthetic pATF promoters. **A**. Circuit design of dCas9 based artificial transcription factor-controlled activation of synthetic promoters (pATFs). Specific *gRNAs* are produced by U6 promoter while the expression of the dCas9-VP64 is under the control of the 35S promoter. Reporter genes are under the control of the synthetic promoter (3 repeats of the *gRNA* followed by minimal 35S promoter to the artificial promoter (*gRNA* binding site) upstream of a specific fluorescence reporter. **B**. Fluorescence microscope image showing Agrobacterium mediated transient expression of YFP, BFP and RFP into *Nicotiana benthamiana* leaves with dCas9-VP64 (bottom panels) and without dCas9-VP64 (upper panels) using three different *gRNAs*. Images were captured using the appropriate filter (Materials and Methods) at same exposure. **C**. Relative integrated density of each fluorescence signal (shown in panel B). Integrated density was measured using image J software and normalized to that of the control (con; - dCAS9-VP64). Error bars: S.D. (n=3, independent replicates). Asterisks indicate statistical significance in a student t-test (P<0.05).

In order to show that the OCS designs could also function in stable transgenic *Arabidopsis thaliana* lines, we assembled the OCS 1-1 and 4-1 circuits (**Supplementary Table 2**; constitutive YFP and luciferase expression, respectively) in an *Agrobacterium* expression vector containing with a kanamycin selectable marker as described previously. These OCS constructs were successfully transformed into *Arabidopsis thaliana* plants (**Fig 4A**). As expected, the OCS 1-1 T_1_ plants exhibited constitutive YFP expression (**Fig 4B**) while the OCS 4-1 plants were imaged (as described in **Methods**) and the constitutive expression of luciferase was confirmed (**Fig 4C, 4D**). Thus, the modular circuits assembled function in two species, as infiltrates in *Nicotiana* and as transgenics in *Arabidopsis*.

**Figure 4.**
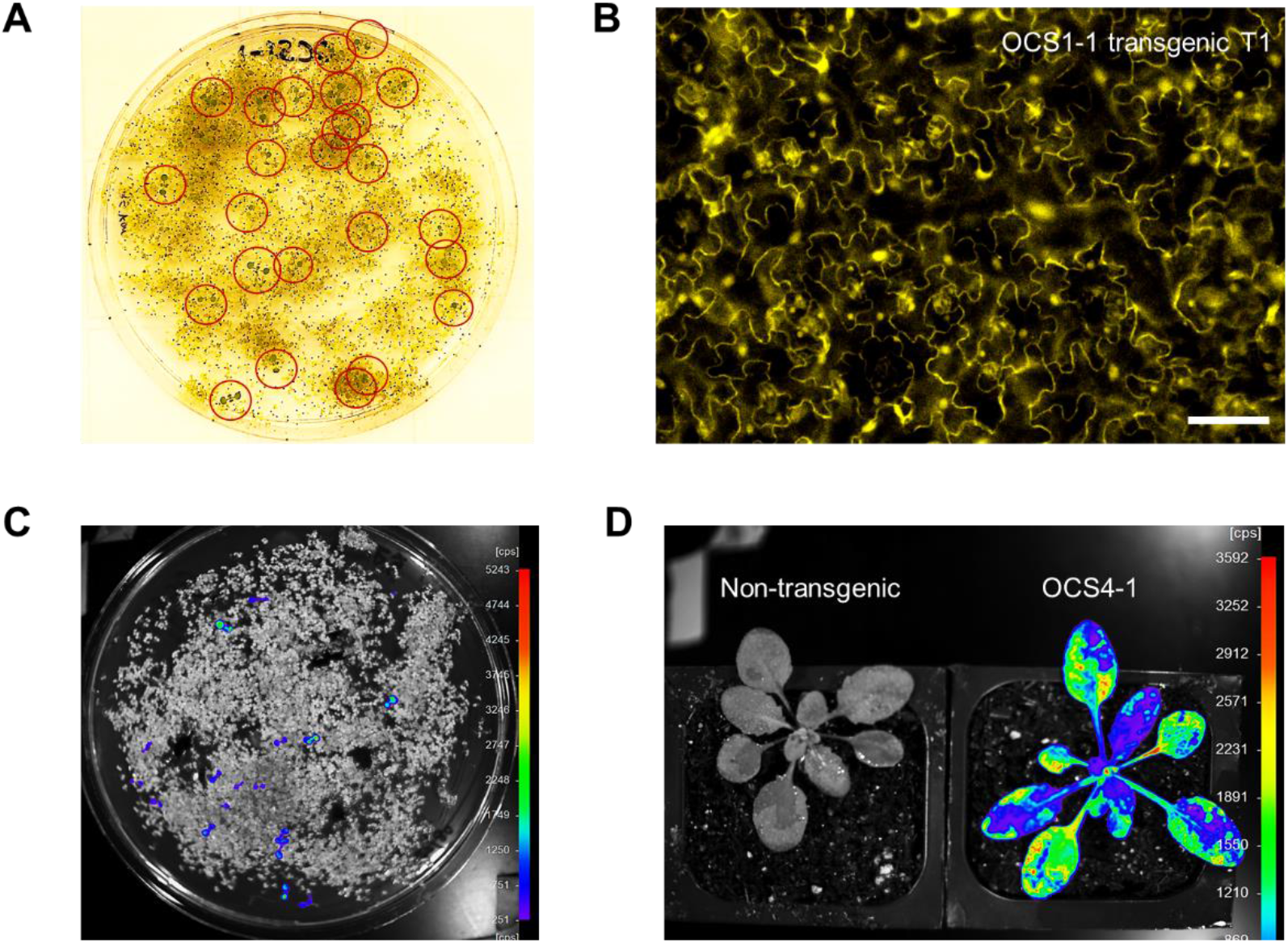
Evaluation of OCS reporter gene expression in transgenic *Arabidopsis* plants. **A**. Image showing Kanamycin selection of the transgenic *Arabidopsis* seedlings on MS media. Seedlings highlighted in the red circle have successfully incorporated OCS circuit. Transformation efficiency is within reasonable ranges (~1%) determined by a simple evaluation of the identified seedlings. **B**. Fluorescence microscope image of *Arabidopsis* transgenic T1 plants containing the constitutive expression of YFP under the OCS control (OCS 1-1). Scale bar: 50 μm **C**. Image showing Kanamycin selection of the transgenic *Arabidopsis* seedlings on MS media using luminescence reporter (OCS4-1) taken using the NightOwl (Methods). **D**. Image of a T1 *Arabidopsis* plant containing OCS4-1 at the rosette stage after spraying the luciferin (Methods) containing OCS4-1. This image, taken at the rosette stage using NightOwl after luciferin spray, shows that the luciferase expression is active throughout the adult plant. A non-transgenic plant on the left was used as a negative control in the luminescence reporter assay.

### Inducible gene expression system via the OCS framework

The ability to precisely regulate the activity of the transgenes/circuit components based on specific input stimuli is a key feature in programmable synthetic circuits (48, 49). In order to enable orthogonal control of induction, we designed gRNA expression cassettes to produce functional gRNAs from inducible Pol II promoters. To prevent nuclear export of gRNAs due to capping and polyadenylation, we used the hammerhead ribozyme (HHR) and Hepatitis Delta Virus (HDV) to cleave the 5’ and the 3’ ends of the gRNA, respectively. This strategy has been previously shown to lead to the expression of functional gRNAs from Pol II promoters, with activity similar to those driven by the Pol III (U6) promoter (50).

To proof the ribozyme processed gRNA constructs, OCS circuits were assembled where gRNAs were either expressed from a U6 promoter (OCS 1-1) or the 35S promoter (OCS 1-5), and could subsequently activate the transcription and expression of reporter genes (YFP) (**Fig 5A**). For both OCS circuits, downstream reporter gene expression was observed, at similar levels (**Fig. 5B**). The specific levels of gRNA obtained in each case were analyzed using qRT-PCR (**Fig 5C and 5D**), and as expected the level of gRNA from the strong Pol II (35S) driven expression was higher than those obtained with the U6 promoter while similar levels of reporter expression were observed for both cases, thus demonstrating that this Pol II driven gRNA expression strategy can be effectively used for OCS activation (**Fig 5E**). For both these constructs the expression of hdCas9 (human codon optimized dCas9) was also confirmed via Western blot analysis (**Fig S2**).

**Figure 5.**
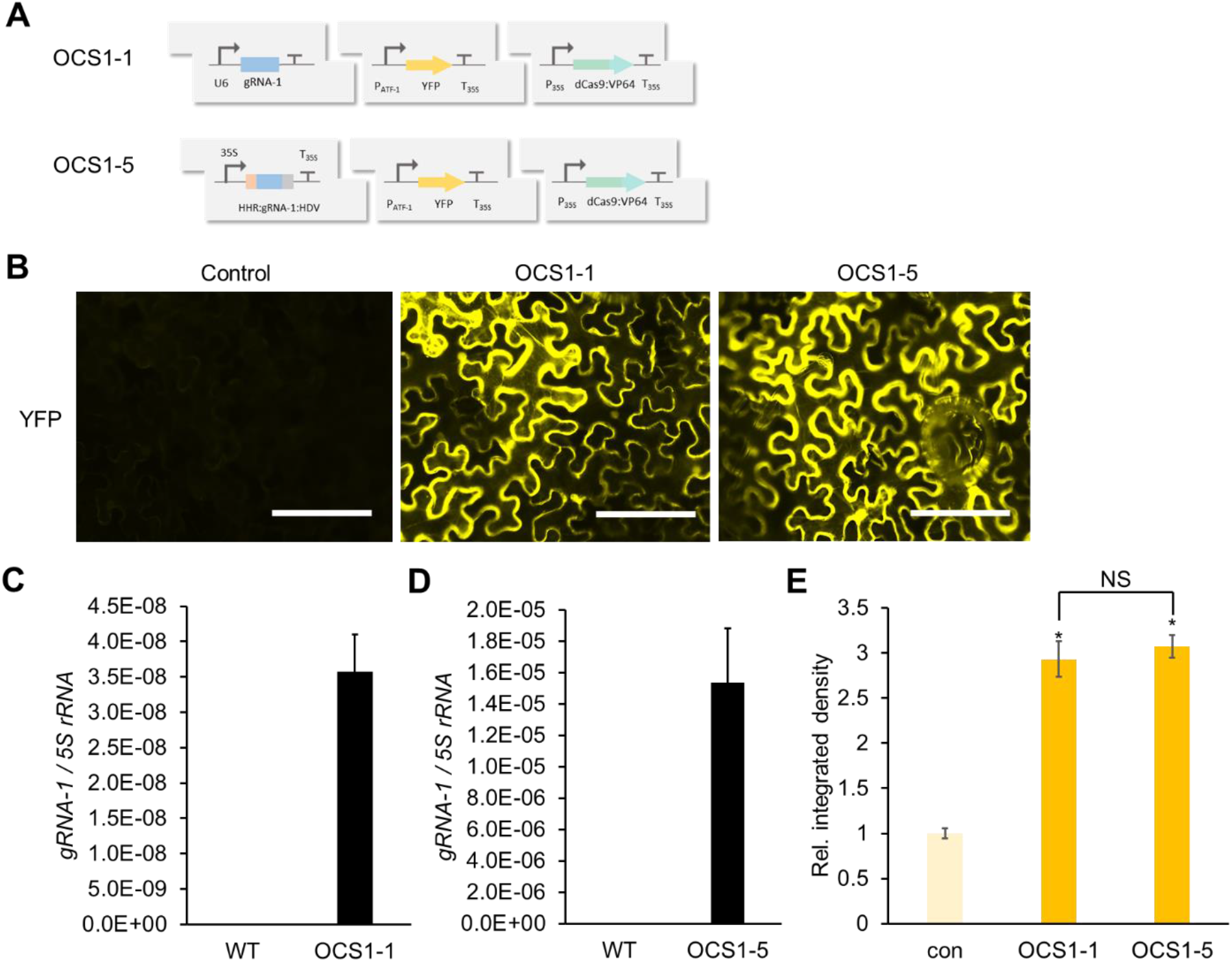
Design and characterization of gRNA expression modules under the control of Pol II promoters. **A**. OCS1-1 circuit generates RNA using U6 (Pol III) promoter while OCS1-5 circuit generates gRNA using 35S (Pol II) promoter flanked by self-cleaving ribozymes – HammerHead (HHR) and Hepatitis Delta Virus (HDV). **B**. Fluorescence microscope images showing *Agrobacterium* mediated transient expression of OCS constructs with two modalities of gRNA expression (OCS1-1 and OCS1-5). Control images were taken without dCAS9-VP64 expression. Scale bars: 200 μm **C** and **D**. Quantification of the *gRNA-1* expression in OCS constructs (OCS 1-1 (C) and OCS 1-5 (D)) using qPCR relative to *5S rRNA*. Error bars : S.D. (n=3, independent replicates) **E**. Relative integrated density of each fluorescence signal (shown in panel B). Integrated density was measured using image J software and normalized to that of the control (con; - dCas9-VP64). Error bars: S.D. (n=3, independent replicates). Asterisks indicate statistical significance in a student t-test (P<0.05). NS: not significant.

In order to demonstrate that the Pol II-driven gRNAs could be used as part of an inducible OCS we used the well-characterized synthetic EBS promoter containing the EIN3 binding (51), and placed YFP under the downstream control of the ATF (via pATF-1) (**Fig 6A**). This circuit (OCS1-9) should be inducible by the volatile organic compound (VOC) ethylene, which is produced from its precursor ACC (1-aminocyclopropane-1-carboxylic acid). Time-dependent expression of YFP is observed in response to 10uM ACC induction (**Fig 6B**). Both the gRNA-1 and YFP expression levels were quantified before and after induction by qRT-PCR, a maximum of 3-fold induction was observed for both cases (**Fig 6C and 6D**). Thus, this demonstrates that the activity from synthetic promoters can be controlled via the selective expression of the corresponding gRNAs.

**Figure 6.**
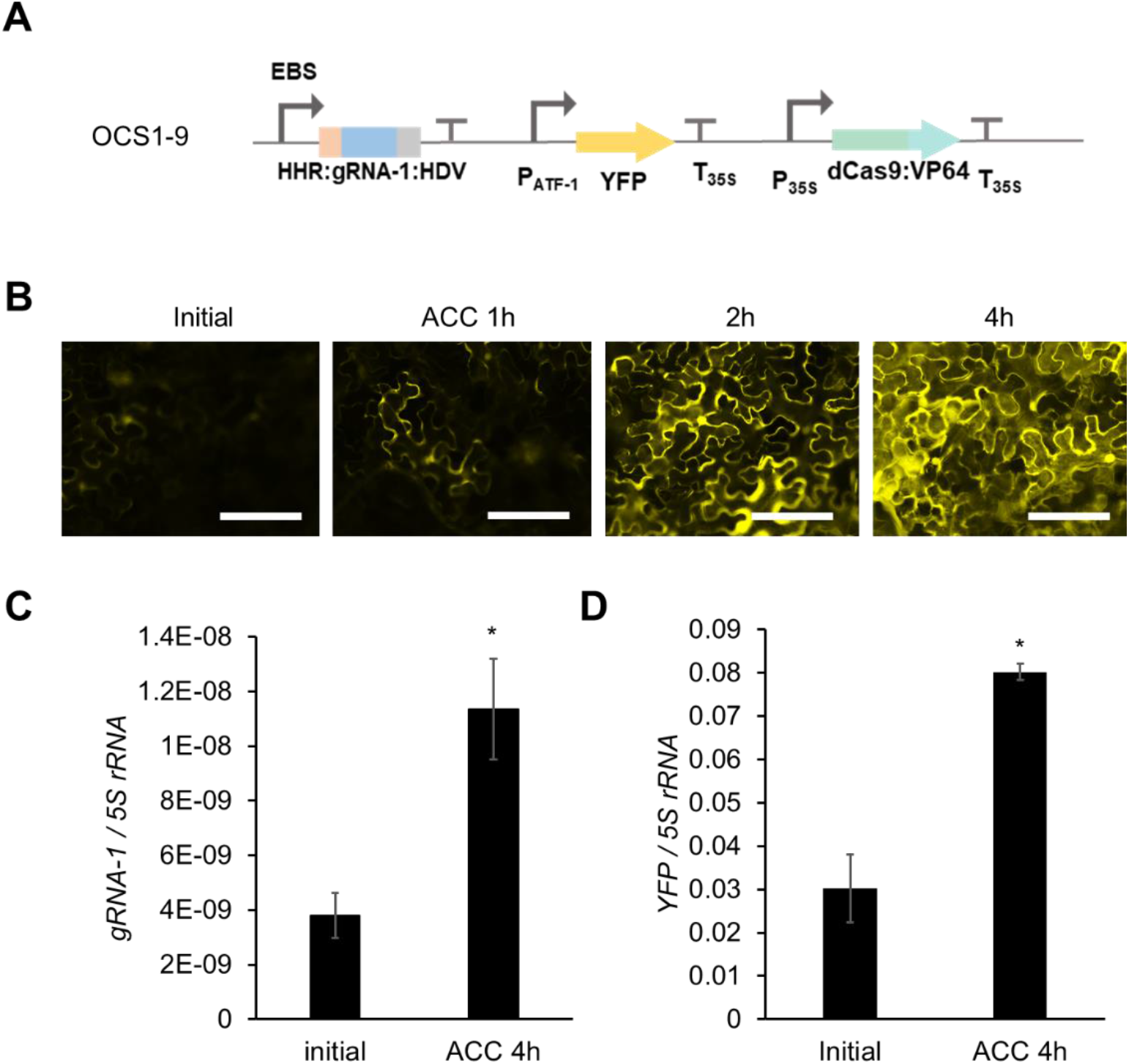
Characterization of an ethylene inducible orthogonal control system. **A**. OCS1-9 circuit (*gRNA-1* is expressed by ethylene inducible EBS promoter) **B**. Time course fluorescence microscope images showing *Agrobacterium* mediated transient expression of OCS1-9 in *Nicotiana benthamiana* leaves after induction with 10μM ACC. Scale bars: 200 μm **C and D**. qPCR quantification of *gRNA-1* (C) and *YFP* (D) expression before and after induction with ACC, where both show similar levels of induction demonstrating that the relative change in *gRNA-1* expression (ethylene induction) results in the differential activation from the pATF-1 promoter. Error bars: S.D. (n=3, independent replicates), Asterisks indicate statistical significance in a student t-test (P<0.05).

### Construction of a panel of mutually orthogonal synthetic promoters

Lack of multiplexed control of transgenes has been a major factor limiting the development of synthetic circuits in plants (5, 6). Multiplexed regulation in turn requires a panel of mutually orthogonal promoters and control elements that can operate simultaneously (5, 6). Our strategy for synthetic promoter design naturally leads to the generation of expression cassettes that are not only orthogonal to the host but are also mutually orthogonal. The degree of orthogonality can be tuned at will via the sequence design of the multiple gRNA components. By simply minimizing homology between gRNAs, we constructed two additional promoters similar to the architecture of pATF-1, in which gRNA binding sites were followed by a minimal 35S promoter (pATF-3 and pATF-4). The orthogonality of these promoters was assayed by assembling expression constructs in which each synthetic promoter controlled the production of a unique fluorescent reporter (pATF-1: YFP, pATF-3: RFP and pATF-4: BFP). The respective gRNAs (gRNA-1, gRNA-3 and gRNA-4) were separately transcribed from a U6 promoter (**Fig 7A**). When expression constructs were infiltrated into *Nicotiana benthamiana*, each of the synthetic promoters was specifically upregulated only when its corresponding gRNA was expressed; no background was detected from the remaining two synthetic promoters. (**Fig 7B and 7C**).

**Figure 7.**
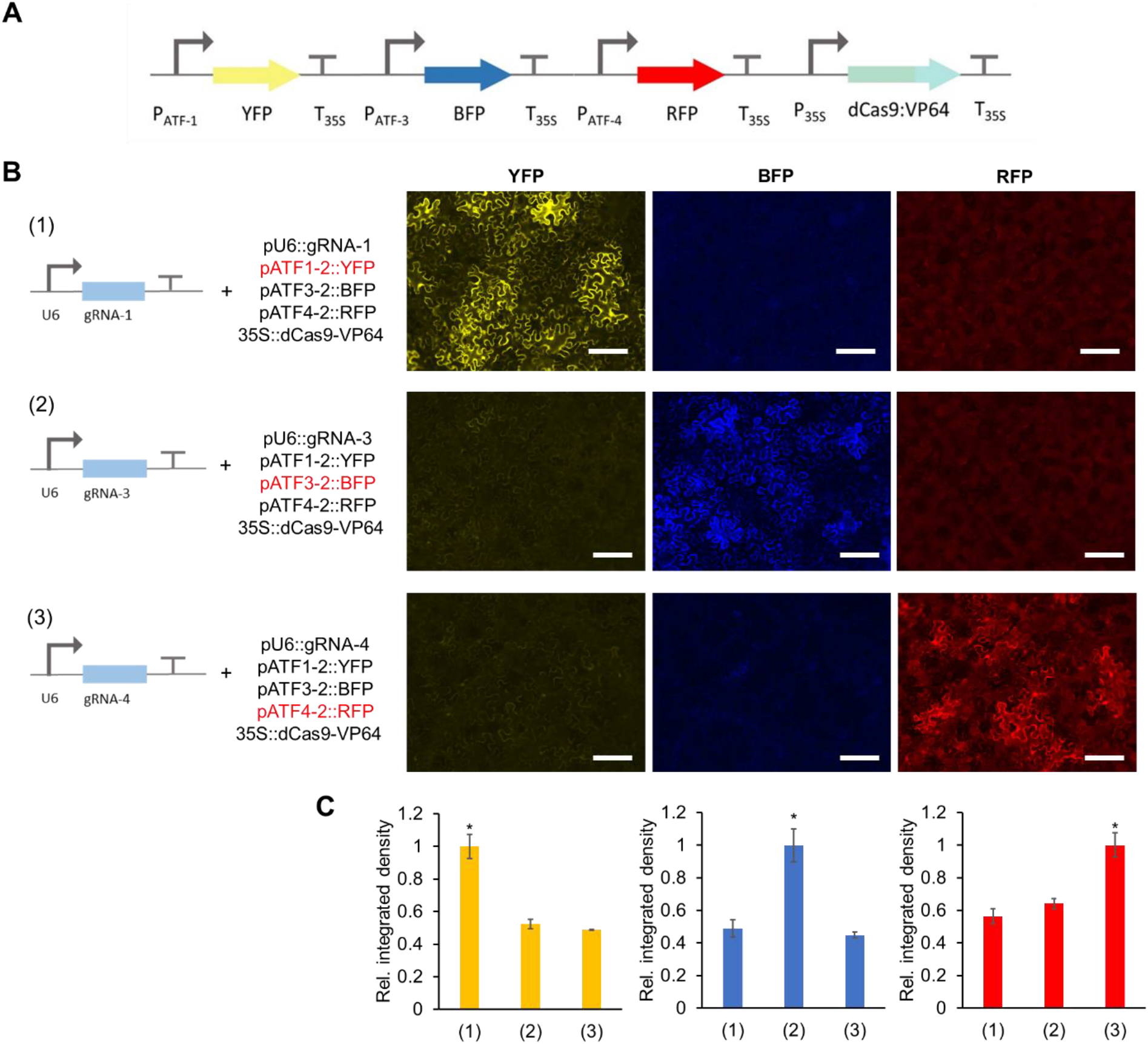
Degree of orthogonality of synthetic promoters. **A**. OCS circuit containing all three synthetic promoters (pATF-1, pATF-3 and pATF-4) driving three different reporter genes namely YFP, BFP and RFP respectively with a single gRNA expressed one at a time under the control of U6 promoter. **B**. Fluorescence microscope images showing *Agrobacterium* mediated transient expression of OCS constructs in *Nicotiana benthamiana* leaves. Scale bars: 200 μm **C**. As observed from the fluorescence images, only the specific gRNA:pATF pair is active, thus demonstrating that the synthetic promoters are mutually orthogonal Relative integrated density of each fluorescence signal (shown in panel B). Integrated density was measured by image J software and normalized to the highest value. Error bars: S.D. (n=3, independent replicates). Asterisks indicate statistical significance in a student t-test (P<0.05).

### Construction of complex ratiometric circuits

Now that we have a suite of mutually orthogonal promoters, we sought to construct simple circuits where the activity of each promoter could be independently controlled. Three separate reporter proteins were used to simultaneously monitor the activity of two promoters: pATF-1 with YFP, while both RFP and BFP were under the control of the pATF-3. By leveraging the designed, orthogonal behavior of these promoters it proved possible to construct a ratiometric circuit wherein the activity of pATF-1, and hence YFP expression, was under the control of ethylene (via ACC), while pATF-3 constitutively drove the expression of RFP and BFP (**Fig 8A**). As expected, the addition of 10uM ACC, induced the expression of YFP from the pATF-1 promoter (3-fold), while the expression of the other reporters remained constant (**Fig 8B and 8C**). The ratiometric response was further validated by qRT-PCR; pATF-1 was induced 3-fold following a similar increase in expression of gRNA-1 while there were no changes observed in the transcription of the other two reporter genes (**Fig 8B and 8C**).The predictable behavior of the designed, artificial control elements in the ratiometric circuit is one of the first examples of complex circuitry to be described in plants, and demonstrates uniquely how natural metabolism and regulatory circuitry can be interfaced with free-standing orthogonal control systems.

**Figure 8.**
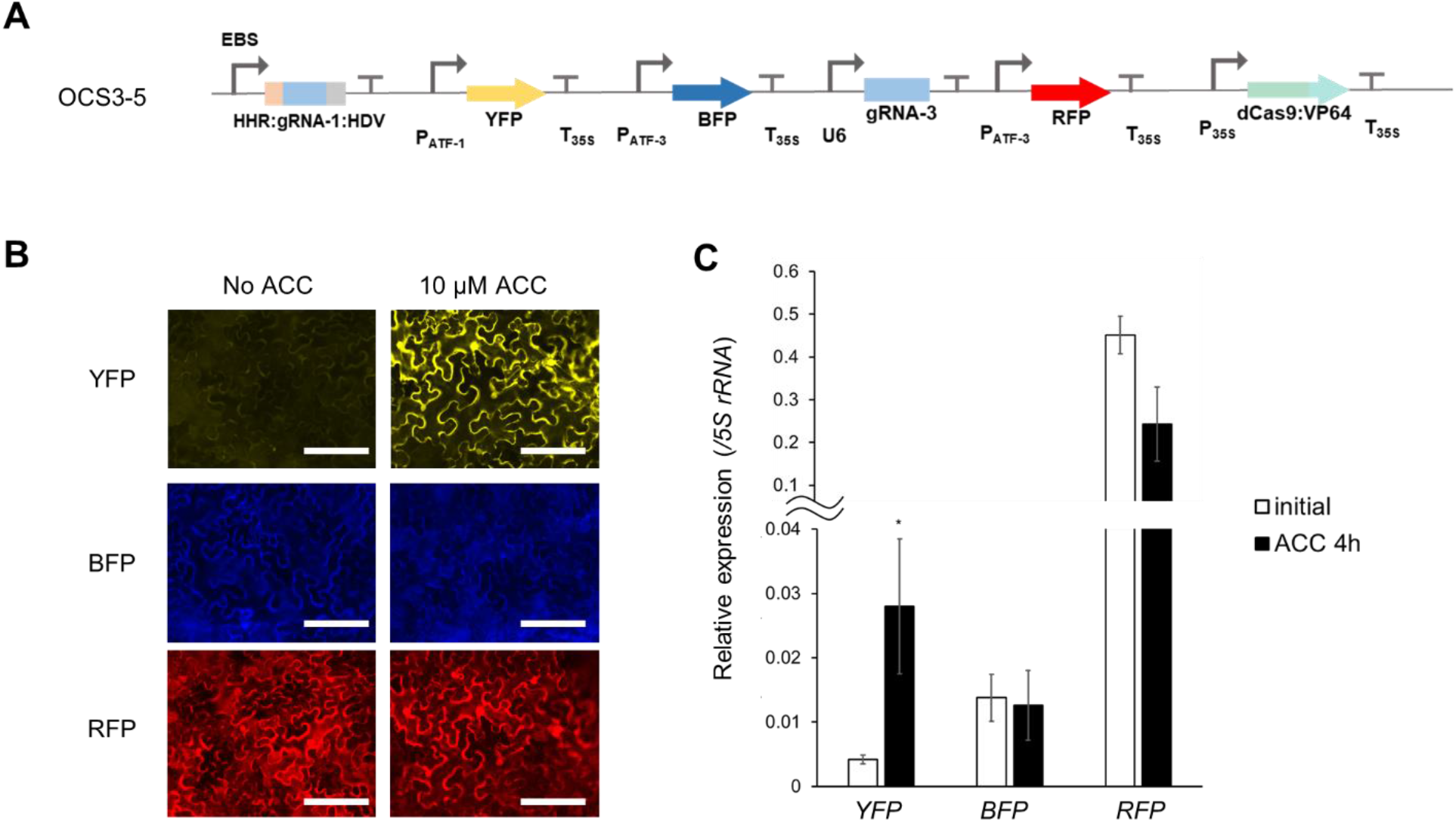
Design and characterization of a ratiometric circuit. **A**. OSC3-5 contains YFP which is inducible by ACC (pATF-1), while BFP and RFP are constitutively expressed under the control of pATF-3 via the constitutive expression of gRNA-3. **B**. Fluorescence microscope images showing *Agrobacterium* mediated transient expression of the ratiometric OCS construct (OCS3-5) in *Nicotiana benthamiana* leaves with or without 10μM ACC. Scale bars: 200 μm **C**. qPCR quantification of *YFP*, *BFP* and *RFP* shows that YFP is induced after the treatment with ACC while the expression of BFP and RFP remains unchanged before or after ACC induction. Error bars: S.D. (n=4, independent replicates). An asterisk indicates statistical significance in a student t-test (P < 0.05).

## Discussion

Transcriptional orthogonality is one of the bedrocks for circuit construction in synthetic biology, and generally serves as the basis for the bottom-up construction of complex circuitry for predictable dynamics (7, 10, 17). For eukaryotes the construction of multiple promoter elements is hindered by the typically complex regulatory sequences that lie upstream and within promoters (52–54).

The design of synthetic eukaryotic promoters has traditionally implemented a common architecture, where a strong transcriptional initiation region is cloned downstream of orthogonal DNA binding operator sequences and the latter serve as landing pads for synthetic transcription factors (23). The engineered transcription factors have typically consisted of DNA binding proteins (i.e., prokaryotic DNA binding proteins like TetR, LacI, LexA and PhIF (55–57)) fused to well characterized transcriptional activation domain like VP64. With the advent of programmable DNA binding proteins like zinc finger proteins and TALEs the repertoire of synthetic promoters greatly increased (23, 24, 58, 59). That said, each new synthetic promoter still requires the construction and characterization of its own unique transcription factor (23, 59, 60).

These bottlenecks can be circumvented by the use of the highly programmable RNA-guided DNA binding protein dCas9 (26). The dCas9 RNP fused to transcription activation domains such as VP64 has been used for the upregulation of endogenous genes in a wide variety of eukaryotic species like yeast, mammalian cells and plants (16, 25, 26, 61). Here, we have used adapted this ‘universal’ transcription factor to control the expression of synthetic and orthogonal promoters without the need of addition of any other factors. Using our modular framework, we were able to quickly design and characterize a panel of mutually orthogonal promoters that could drive the production of a variety of outputs, singly and in parallel, including different fluorescent proteins (GFP, BFP, RFP and YFP) and luciferase.

The activities of dCas9 based transcription factors can potentially be controlled by simply regulating the expression of their corresponding gRNAs (16, 17), enabling the coupling of natural and synthetic transcription units, and thus natural and overlaid metabolic responses. Here we have effectively used this strategy to couple ethylene sensing (via known EIN3 binding sites) to synthetic (pATF) promoters. Moreover, by changing the number and arrangement of gRNA binding sites synthetic promoters with different levels of activation can be generated, providing further opportunities for design (62). While it has been previously shown that a panel of minimal plant promoters can be used with natural DNA binding sequences for modulating promoter strengths (20), the addition of completely artificial, synthetic promoters as control elements should create opportunities for increasing the specificity and strengths of engineered promoters.

Since our strategy for designing synthetic promoters is generalizable it is likely that even more complex circuits can be built by simply incorporating other transcription factor binding sites, or by changing the regulatory ‘headpiece’ on the dCas9 element (for example, to a repressor), (63–65).

The stabilities of genetic circuitry in plants can be greatly modified by silencing and recombination, amongst other mechanisms (40, 41, 43). In this regard, the artificial promoter elements that we generate can potentially be crafted to avoid repetition (20), and thus to better avoid silencing and recombination. As viable artificial promoter sequences continue to accumulate, they can be compared and contrasted to identify those that are least vulnerable to modification over time. The facile addition of new parts to the standardize toolkit architecture, particularly terminators, will further increase opportunities to avoid repetition in ways that again go well beyond what is possible by relying on just a few well-characterized endogenous elements alone.

The implementation of orthogonal control systems in plants can be used to limit cross-talk between natural and overlaid regulatory elements, allowing more precise response to a variety of inputs, from VOCs to hormones to temperature, water, and nutrients. The use of orthogonal control systems to enable more precise responses to pathogenesis is especially intriguing given the presence of R genes that are specifically responsive to individual pathogens (effector triggered immunity, ETI) (66). The architecture we have developed is fully generalizable, and can potentially be expanded to non-model plants and other eukaryotic species such as yeast and mammalian cells by the use of appropriate transcription initiation regions under the control of similar gRNA sequences binding sites (67).

## Materials and Methods

### Plasmid design and construction

The plant expression vector was generated using the plasmid pICH86966 (Addgene#48075) as the backbone. The lacZ expression cassette was replaced with the GFP dropout sequence (**Supplementary Table 2**) to make the plasmid compatible with YTK architecture design. All parts described in **Supplementary Table 1**, were cloned into the backbone pYTK001 (Addgene #65108). For the individual transcriptional units, the backbone used was pYTK095 (Addgene #65202) along with the appropriate connector sequences described in **Supplementary Table 3**. For the design of orthogonal gRNAs, random 20-mers were generated that had a GC content of ~50%, and that were at least 5 nucleotides away from all sequences in the *Nicotiana* and *Arabidopsis* genomes. All oligonucleotides and gblocks were obtained from Integrated DNA Technologies (IDT) unless otherwise stated.

For the construction of each genetic element namely promoters, coding sequences and terminators, first they were checked for restriction sites for the following enzymes – BsmBI, BsaI, NotI and DraIII. The restriction sites in the coding sequences were removed by the use of synonymous codons while the other elements did not contain any of these restriction sites. The complete list of parts and constructs are provided in **Supplementary Table 1**. The part plasmids were cloned into a common vector where each genetic element is flanked by Bsa1 restriction sites followed by appropriate overhangs (**Supplementary Table 1**). For the assembly of both single TU or multi-TU, the following procedure was used: 10 fmol of backbone plasmid and 20 fmol of parts/TUs were used in a 10uL reaction with 1ul of 10x T4 ligase buffer along with 100 units of BsaI-v2 (single TU) or Esp3I (multi-TU or parts) and 100 units of T7 DNA ligase. The cycling protocol used is: 24 cycles of 3 min at 37°C (for digestion) and 5 min at 16°C (for ligation) followed by a final digestion step at 37°C for 30min and the enzymes were heat inactivated 80°C for 20 min. All constructs were transformed into DH10B cells, grown at 37°C using standard chemical transformation procedures. The colonies that lack fluorescence were inoculated and plasmids were extracted using Qiagen Miniprep kit according to the manufacturer’s instructions Plasmids were maintained as the following antibiotics kanamycin (50ug/mL), chloramphenicol (34ug/mL) and carbenicillin (100ug/mL) wherever required. The plasmids were sequence verified by Sanger sequencing (UT Austin Genomic Sequencing and Analysis Facility). The correct constructs were then transformed into *Agrobacterium tumefaciens* strain GV3101 (resistant to Gentamycin and Rifampicin) and used either for transient expression in *Nicotiana benthamiana* or to generate stable lines in *Arabidopsis thaliana*. The following enzymes were used for the assemblies – BsaI-v2 (NEB #R3733S), Esp3I (NEB #R0734S) and T7 DNA ligase (NEB #M0318S).

### Plant material, bacterial infiltration

*Nicotiana benthamiana* and *Arabidopsis thaliana* plants were grown in soil at 22°C, and 16 hr light period. For transient expression, three weeks old plants were syringe-infiltrated with *Agrobacterium tumefaciens* strain GV3101 (OD_600_ = 0.5) and leaves were imaged under Olympus BX53 Digital Fluorescence Microscope or harvested for RNA and/or protein analysis. To create stable transformation in Arabidopsis, floral dip method (68) was used. T_1_ plants were selected on half MS Kanamycin (50μg/ml) plates and the selected T1 plants were analyzed using an Olympus BX53 Digital Fluorescence Microscope and a NightOwl imager for YFP expression and luciferase expression, respectively. For circuits that constitutively expressed YFP (OCS1-1) and luciferase (OCS4-1) no other obvious phenotypic differences were observed across numerous individual plants.

### RNA extraction and qRT-PCR

RNA was extracted using TRIzol reagent (Ambion). 1μg total RNA was used to synthesize cDNA. After DNaseI treatment to remove any DNA contamination, random primer mix (NEB #S1330S) and M-MLV Reverse transcriptase (Invitrogen #28025-013) were used for first strand synthesis. qRT-PCR was used to quantify the RNA prepared from transient expression experiments. AzuraQuant qPCR Master Mix (Azura Genomics) was used with initial incubation at 95 °C for 2 min followed by 40 cycles of 95 °C for 10 sec and 60 °C for 30sec. Level of target RNA was calculated from the difference of threshold cycle (Ct) values between reference (*5S rRNA*) and target gene using at least three independent replicates

### ACC treatment

To check the induction of reporter in response to ACC in the plasmids containing pEBS∷YFP/RFP/BFP, *Nicotiana benthamiana* leaves were infiltrated with Agrobacterium; after three days post infiltration, leaf discs were cut using cork borer and incubated in either 0μM or 10μM ACC for four hours. Fluorescence microscopy was used to check YFP expression after induction.

#### Fluorescence and Luminescence imaging

Fluorescence microscope images after *Agrobacterium* mediated transient expression of YFP, BFP, RFP and GFP in *Nicotiana benthamiana* leaves were taken using an Olympus BX53 Digital Fluorescence Microscope. For this purpose, leaf discs were cut using cork borer from the area which was infiltrated. Images were taken using either 10X objective lens using the default filters for YFP (500/535nm), BFP (385/448nm), and RFP (560/630nm). The UV filter (350/460nm) was used to take GFP images. The exposure and gain setting were kept constant for each filter within each experiment to compare multiple leaf discs (3 to 6). In all the experiments a leaf disc from a leaf which was not infiltrated with Agrobacterium was used as a negative control in order to account for background fluorescence. All experiments were performed at least three times independently as indicated in the Results.

Expression of luciferase was detected using NightOwl II LB 983 *in vivo* imaging system (https://www.berthold.com/en/bioanalytic/products/in-vivo-imaging-systems/nightowl-lb983/). Leaves/plants were sprayed with 100μM D-luciferin, Potassium salt (GoldBio #LUCK-300). After 5 min of incubation, images were taken in the NightOwl II LB 983. Images were captured with a backlit NightOWL LB 983 NC 100 CCD camera. Photons emitted from luciferase were collected and integrated for a 2 min period. A pseudocolor luminescent image from blue (least intense) to red (most intense), representing the distribution of the detected photons emitted from active luciferase was generated using Indigo software (Berthold Technologies).

### Western blot

Total protein was extracted using urea-based denaturing buffer (100 mM NaH2PO4, 8 M urea, and 10 mM Tris-HCl, pH 8.0) and used for immunoblot analysis to check the expression. The proteins were fractionated by 8% SDS-PAGE gel and transferred to a polyvinylidene difluoride (PVDF) membrane using a transfer apparatus according to the manufacturer’s protocols (Bio-Rad). The membrane was treated with 5% nonfat milk in PBS-T for 10 min for blocking, and then incubated with Cas9 antibody (Santa cruz, 7A9-3A3, 1:500) at 4 °C for overnight. After incubation, the membrane was washed three times for 5 min and incubated with horseradish peroxidase-conjugated anti-mouse (1:10000) for 2 h. The Blot was washed with PBS-T three times and detected with the ECL system (Thermo scientific, lot# SE251206).

## Declarations

### Ethics approval and consent to participate

Not Applicable

### Consent for publication

Not Applicable

### Availability of data and materials

### Competing interests

The authors declare no competing interests.

### Funding

This work was supported by the Defense Advanced Research Projects Agency (DARPA) agreement HR00111820048 to AE and SS. The content of the information does not necessarily reflect the position or the policy of the Government, and no official endorsement should be inferred. This work was also supported by Welch Foundation grant (F-1654) to ADE.

### Authors’ contributions

SK, SS and AE conceived of the project. SK designed the framework and the basic elements of OCS with input from EG, JG and SS. SK and YB assembled all constructs. YB, NR and JK performed all the testing in Nicotiana with input from SS. All authors contributed with the preparation of figures. SK, YB, JK, SS and AE wrote the manuscript with input from all authors.

## Acknowledgments

We would also like to thank the Qiao lab (UT Austin) for providing details regarding the ethylene induction of the OCS constructs.

## Supplementary Information includes

**Fig S1**: Workflow describing the assembly of single and multiple transcriptional units (TUs) in a plant expression vector; **Fig S2**: Western blot to analyze the expression of dCas9:VP64 in OCS constructs – OCS1-1 and OCS 1-5

**Table S1**: List of all genetic parts used for the construction of OCS constructs

**Table S2**: List of all OCS constructs

**Table S3**: List of all Addgene plasmids used in this work

**Full OCS plasmid maps**

## Notes

### Competing Interest Statement

The authors have declared no competing interest.

