## Supplementary information for "Orthogonal control of gene expression in plants using synthetic promoters and CRISPR-based transcription factors"

**Figure S1**: Workflow describing the assembly of single and multiple transcriptional units (TUs) in a plant expression vector

**Figure S2**: Western blot to analyze the expression of dCas9:VP64 in OCS constructs – OCS1-1 and OCS 1-5

**Table S1**: List of all genetic parts used for the construction of OCS constructs

**Table S2**: List of all OCS constructs

**Table S3**: List of all Addgene plasmids used in this work

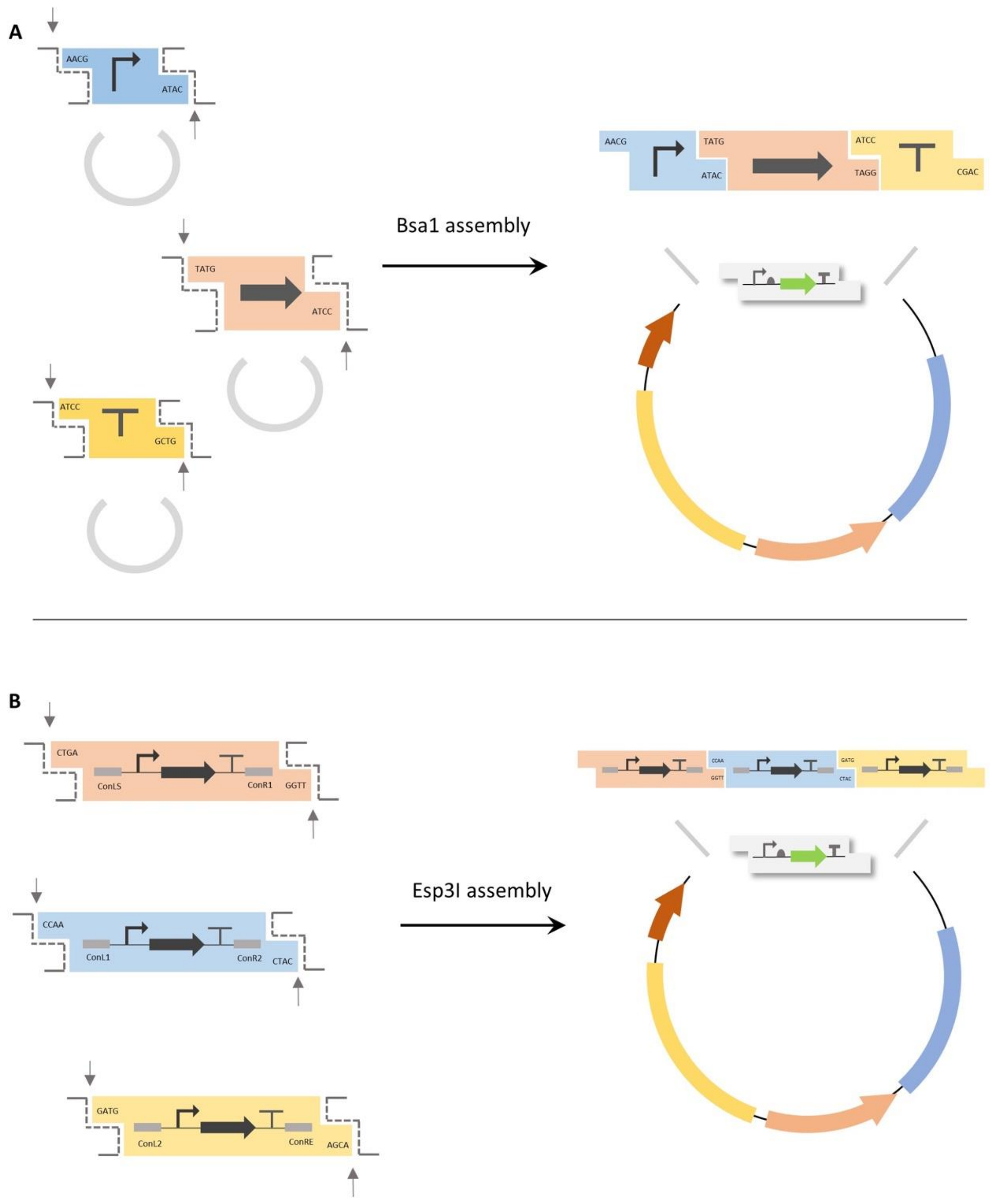


**Supplementary figure 1**: Schematic demonstrating the assembly of single (A) or multiple (B) transcriptional units into a plant expression vector. A single transcription unit consists of a promoter, gene and a terminator parts while for multiple transcriptional units, each TU is flanked by appropriate connector sequences. The arrows depict the restriction sites for Bsa1 (A) and Esp3I (B). B) Schematic showing the assembly of multiple TUs into plant expression vector where each TU is encoded in separate plasmids flanked by appropriate connector sequences.

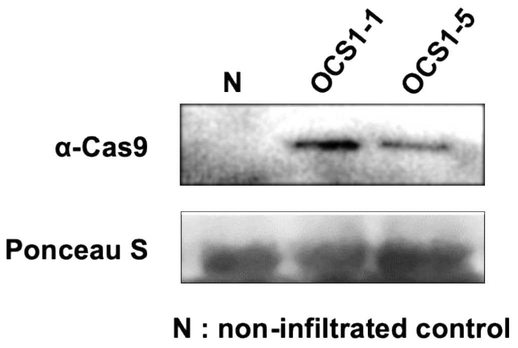


**Supplementary figure 2**: The expression of hdCas9 for both OCS 1-1 and OCS 1-5 was confirmed via Western blot analysis (see Methods).

**Table S1: List of genetic elements used.**

Highlighted regions in red indicate BsaI recognition sites and the corresponding overhangs are highlighted in blue.

**S1.1 Promoters**

| Name | Description | Sequence |
|---|---|---|
| P1 | 35S promoter | GGTCTCAAACGCGTCAACATGGTGGAGCACGACAC<br>TCTGGTCTACTCCAAAAATGTCAAAGATACAGTCTCA<br>GAAGATCAAAGGGCTATTGAGACTTTTCAACAAAGG<br>ATAATTTCGGGAAACCTCCTCGGATTCCATTGCCCA<br>GCTATCTGTCACTTCATCGAAAGGACAGTAGAAAAG<br>GAAGGTGGCTCCTACAAATGCCATCATTGCGATAAA<br>GGAAAGGCTATCATTCAAGATCTCTCTGCCGACAGT<br>GGTCCCAAAGATGGACCCCCACCCACGAGGAGCAT<br>CGTGGAAAAAGAAGAGGTTCCAACCACGTCTACAAA<br>GCAAGTGGATTGATGTGACATCTCCACTGACGTAAG<br>GGATGACGCACAATCCCACTATCCTTCGCAAGACCC<br>TTCCTCTATATAAGGAAGTTCATTTCATTTGGAGAGG<br>ACACGCGTTTATTTACAAGAGCGTACGGTTCAATCC<br>CTGCCTCCCCTGTAAAACTACCCTTTGAAAACCTCTC<br>TTTCTTAATCTTTTCTTTGTAATTCCAGATCTATGTGA<br>GACC |
| P2 | Mas promoter | GGTCTCAAACGCGGAGATTTTTCAAATCAGTGCGCA<br>AGACGTGACGTAAGTATCCGAGTCAGTTTTTATTTTT<br>CTACTAATTTGGTCGTTTATTTCGGCGTGTAGGACAT<br>GGCAACCGGGCCTGAATTTCGCGGGTATTCTGTTTC<br>TATTCCAACTTTTTCTTGATCCGCAGCCATTAACGAC<br>TTTTGAATAGATACGCTGACACGCCAAGCCTCGCTA<br>GTCAAAAGTGTACCAAACAACGCTTTACAGCAAGAA<br>CGGAATGCGCGTGACGCTCGCGGTGACGCCATTTC<br>GCCTTTTCAGAAATGGATAAATAGCCTTGCTTCCTAT<br>TATATCTTCCCAAATTACCAATACATTACACTAGCAT<br>CTGAATTTCATAACCAATCTCGATACACCAAATCGAG<br>ATCTATGTGAGACC |
| P4 | EBS promoter | GGTCTCAAACGCATTACGAATTCCCGGGATCCTCA<br>TGATCAAAGGGGGGATGCACTATTTAAGATCCTCAT<br>GATCAAAGGGGGGATGCACTATTTAAGATCCTCATG<br>ATCAAAGGGGGGATGCACTATTTAAGATCCTCATGA<br>TCAAAGGGGGGATGCACTATTTAAGATCCTCATGAT<br>CAAAGGGGGGATGCACTATTTAAGATCCTTCGCAAG<br>ACCCTTCCTCTATATAAGGAAGTTCATTTCATTTGGA<br>GAGGACAGGGTATCAAGCTTGGCACTGGCCGTCGT<br>TTTACAACGTCGTGACTGGGAAAACCCTGGCGTTAC |

| | | |
|---|---|---|
| | | CCAACTTAATCGCCTTGCAGCACATCCCCCTTTCGC<br>CAGCTGGCGTAATAGCGAAGAGGCCCGCACCGATC<br>GCCCTTCCCAACAGTTGCGCAGCCTGAAGCCTAGG<br>GAGGAGTCCACTCTATGTGAGACC |
| pATF-1 | Artificial promoter 1 (3 gRNA repeats) | GGTCTCAAACGCCAAACCACAATTTGCACACCCTGG<br>CATCTCAAACCACAATTTGCACACCCTGGCATCTCA<br>AACCACAATTTGCACACCCTGGCATCTCCGCAAGAC<br>CCTTCCTCTATATAAGGAAGTTCATTTCATTTGGAGA<br>GGACACGCGTTTATTTACAAGAGCGTACGGTTCAAT<br>CCCTGCCTCCCCTGTAAAACTACCCTTTGAAAACCT<br>CTCTTTCTTAATCTTTTCTTTGTAATTCCAGATCTATG<br>TGAGACC |
| pATF-3 | Artificial promoter 3 (3 gRNA repeats) | GGTCTCAAACGCCTGACTCACAGTCCTATCGAGTGG<br>CATCTCTGACTCACAGTCCTATCGAGTGGCATCTCT<br>GACTCACAGTCCTATCGAGTGGCATCTCCGCAAGAC<br>CCTTCCTCTATATAAGGAAGTTCATTTCATTTGGAGA<br>GGACACGCGTTTATTTACAAGAGCGTACGGTTCAAT<br>CCCTGCCTCCCCTGTAAAACTACCCTTTGAAAACCT<br>CTCTTTCTTAATCTTTTCTTTGTAATTCCAGATCTATG<br>TGAGACC |
| pATF-4 | Artificial promoter 4 (3 gRNA repeats) | GGTCTCAAACGCCATTGCGCACCATTCCACTAGTGG<br>CATCTCATTGCGCACCATTCCACTAGTGGCATCTCA<br>TTGCGCACCATTCCACTAGTGGCATCTCCGCAAGAC<br>CCTTCCTCTATATAAGGAAGTTCATTTCATTTGGAGA<br>GGACACGCGTTTATTTACAAGAGCGTACGGTTCAAT<br>CCCTGCCTCCCCTGTAAAACTACCCTTTGAAAACCT<br>CTCTTTCTTAATCTTTTCTTTGTAATTCCAGATCTATG<br>TGAGACC |

#### S1.2 Terminators

| Name | Description | Sequence |
|---|---|---|
| T1 | 35S Terminator | GGTCTCAATCCTAACTCGAGCTCTAGCTAGAGTCGA<br>TCGACAAGCTCGAGTTTCTCCATAATAATGTGTGAGT<br>AGTTCCCAGATAAGGGAATTAGGGTTCCTATAGGGT<br>TTCGCTCATGTGTTGAGCATATAAGAAACCCTTAGTA<br>TGTATTTGTATTTGTAAAATACTTCTATCAATAAAATT<br>TCTAATTCCTAAAACCAAAATCCAGTACTAAAATCCA<br>GATGCTGTGAGACC |

#### S1.3 Coding sequences

| Name | Description | Sequence |
|---|---|---|

| | | |
|---|---|---|
| G1 | YFP | GGTCTCATATGGTATCCAAAGGAGAAGAATTGTTTA<br>CAGGGGTAGTCCCCATATTGGTCGAGCTCGATGGG<br>GATGTAAACGGGCACAAGTTCTCAGTGTCTGGCGAA<br>GGCGAAGGGGACGCCACATACGGAAAGTTGACTCT<br>GAAGTTCATCTGCACAACCGGTAAACTACCCGTACC<br>CTGGCCAACACTGGTGACGACCTTTGGGTACGGGC<br>TACAATGTTTCGCTCGATATCCTGACCACATGAAGC<br>AACACGATTTCTTCAAAAGCGCCATGCCTGAGGGGT<br>ATGTACAGGAGCGTACCATATTTTTCAAAGACGATG<br>GAAACTATAAGACCCGAGCTGAAGTGAAATTTGAAG<br>GAGATACGTTAGTCAATCGAATAGAACTAAAGGGAA<br>TTGATTTCAAGGAGGATGGGAATATTCTGGGCCATA<br>AGCTGGAGTACAATTACAATAGTCACAACGTCTATAT<br>AATGGCAGACAAGCAGAAGAATGGAATAAAGGTTAA<br>TTTCAAGATAAGGCATAACATTGAAGACGGAAGCGT<br>TCAGCTAGCCGATCATTACCAACAGAATACACCGAT<br>AGGTGATGGACCCGTCCTGCTCCCCGACAACCATTA<br>CTTGTCTTACCAGAGTGCACTTTCTAAAGATCCAAAT<br>GAGAAAAGAGATCATATGGTTTTATTAGAATTTGTTA<br>CCGCAGCCGGCATAACTTTGGGAATGGACGAACTG<br>TACAAATGAGGATCCTGAGACC |
| G2 | RFP | GGTCTCATATGAGCGAGCTGATTAAGGAGAACATGC<br>ACATGAAGCTGTACATGGAGGGCACCGTGAACAAC<br>CACCACTTCAAGTGCACATCCGAGGGCGAAGGCAA<br>GCCCTACGAGGGCACCCAGACCATGAGAATCAAGG<br>TGGTCGAGGGCGGCCCTCTCCCCTTCGCCTTCGAC<br>ATCCTGGCTACCAGCTTCATGTACGGCAGCAGAACC<br>TTCATCAACCACACCCAGGGCATCCCCGACTTCTTT<br>AAGCAGTCCTTCCCTGAGGGCTTCACATGGGAGAG<br>AGTCACCACATACGAAGACGGGGGCGTGCTGACCG<br>CTACCCAGGACACCAGCCTCCAGGACGGCTGCCTC<br>ATCTACAACGTCAAGATCAGAGGGGTGAACTTCCCA<br>TCCAACGGCCCTGTGATGCAGAAGAAAACACTCGG<br>CTGGGAGGCCAACACCGAGATGCTGTACCCCGCTG<br>ACGGCGGCCTGGAAGGCAGAAGCGACATGGCCCTG<br>AAGCTCGTGGGCGGGGGCCACCTGATCTGCAACTT<br>CAAGACCACATACAGATCCAAGAAACCCGCTAAGAA<br>CCTCAAGATGCCCGGCGTCTACTATGTGGACCACAG<br>ACTGGAAAGAATCAAGGAGGCCGACAAAGAAACTTA<br>CGTCGAGCAGCATGAGGTTGCTGTGGCCAGATACT<br>GCGACCTCCCTAGCAAACTGGGGCACAAGTGAGGA<br>TCCTGAGACC |

| | | |
|---|---|---|
| G10 | BFP | GGTCTCATATGAGCGAAGAACTAATCAAGGAAAATATGCACATGAAACTCTACATGGAGGGTACGGTTGACAATCATCATTTCAAATGTACCGCAGAGGGCGAAGGTAAACCCTATGAGGGCACCCAGACTATGCGAATCAAGGTTGTGGAGGGCGGGCCATTGCCCTTCGCTTTTGACATCTTAGCTACTAGTTTCTTATATGGGAGCAAGACCTTTATAGATCACACACAGGGTATCCCGGACTTCTTTAAACAGAGCTTTCCAGAAGGGTTCACCTGGGAAAGGGTGACAACCTATGAAGATGGCGGAGTGCTTACAGCGACACAAGACACCTCCCTACAGGACGGCACCCTAATATATAATGTCAAGATTCGTGGCGTAGATTTCCTCAGCAATGGCCCTGTTATGCAGAAGAAGACACTTGGATGGGAGGCTTTCACTGAAACCCTCTACCCGGCGGATGGAGGCTTAGAGGGGAGAAACGATATGGCGTTAAAGCTGGTCGGGGGATCACACTTGATCGCGCATGCAAAAACTACGTACAGGTCCAAGAAACCAGCAAAGAATCTCAAGATGCCAGGTGTATACTATGTGGATTACCGACTCGAGCGTATTAAGGAAGCAAACGACGAAACGTACGTCGAACAACACGAAGTTGCAGTAGCAAGGTATAGCGACCTTCCCTCCAAGCTAGGACATAAGCTGAATGGGAGCGGATAAGGATCCTGAGACC |
| G17 | Luciferase | GGTCTCATATGGAAGATGCAAAGAATATCAAAAAAGGCCCAGCGCCCTTCTACCCATTAGAAGATGGAACCGCAGGAGAGCAACTTCACAAGGCGATGAAACGATATGCTCTTGTCCCGGGAACCATCGCTTTTACGGACGCACACATAGAGGTTAACATTACCTATGCGGAATATTTTGAAATGTCAGTCAGATTAGCAGAAGCAATGAAACGTTATGGGCTCAACACTAACCATCGTATTGTTGTATGTAGCGAAAACAGCCTGCAGTTTTTTATGCCGGTGCTCGGTGCGCTGTTCATCGGTGTAGCGGTTGCTCCCGCAAACGACATTTACAACGAGAGAGAACTGCTTAACAGCATGAATATCAGCCAACCGACCGTCGTGTTTGTCTCAAAAAAGGGACTACAAAAAATTCTAAATGTCCAAAAGAAGTTACCTATTATCCAGAAAATTATTATTATGGATAGCAAGACGGATTATCAAGGATTCCAATCTATGTACACATTTGTTACGAGCCACTTACCTCCAGGTTTTAACGAATATGATTTTGTGCCTGAGAGCTTTGATCGAGATAAGACCATCGCGTTAATTATGAATAGTTCCGGCTCTACGGGGCTCCCAAAGGGAGTCGCACTACCACATCGAACTGCGTGCGTTAGATTTTCACATGCCAGAGATCCTATCTTCGGGAATCAGATTATTCCGGACACTGCAATACTGAGTGTGGTTCCGTTTCATCACGGGTTCGGGATGTTCACGACACTCGGTTACCTCATATGCGGATTTCGTGTGGTGCTGATGTATAGGTTTGAAGAAGAGTTGTTCCTAAGATCCTTGCAGGATTACAAAATTCAGTCCGCCCTGTTAGTTCCTACCTTATTTTCCTTTTTCGCCAAGTCAACGTT |

| | | |
|---|---|---|
| | | AATTGATAAATATGACCTATCCAACCTCCACGAAATT<br>GCCAGTGGTGGGGCCCCCTTGTCCAAAGAAGTTGG<br>TGAGGCAGTCGCTAAGAGGTTCCACCTGCCGGGTA<br>TCCGTCAAGGCTACGGACTTACCGAAACAACCTCCG<br>CTATTCTTATTACACCTGAAGGCGATGATAAGCCGG<br>GTGCTGTCGGTAAGGTGGTGCCTTTTTTTGAGGCGA<br>AGGTAGTGGACTTAGATACTGGCAAGACGCTCGGA<br>GTTAATCAACGAGGCGAGCTCTGTGTCCGTGGTCCC<br>ATGATAATGAGCGGATACGTCAACAATCCTGAGGCA<br>ACCAACGCATTAATTGATAAAGACGGCTGGTTGCAT<br>AGCGGGGATATCGCGTATTGGGATGAGGACGAACA<br>CTTCTTTATAGTCGATAGGTTAAAGTCACTTATTAAG<br>TATAAAGGTTATCAAGTCGCTCCCGCCGAACTGGAG<br>AGTATTCTTCTTCAGCATCCGAATATCTTTGACGCCG<br>GTGTTGCAGGTCTTCCTGACGACGACGCCGGAGAA<br>CTACCTGCAGCGGTCGTGGTGCTCGAACATGGAAA<br>GACAATGACCGAGAAGGAGATTGTAGATTACGTAGC<br>TTCACAGGTCACCACCGCTAAAAAACTAAGAGGTGG<br>TGTTGTCTTCGTAGATGAAGTGCCTAAAGGACTTAC<br>CGGTAAACTCGACGCCAGGAAGATAAGAGAAATCCT<br>GATCAAAGCGAAGAAGGGTGGTAAGTCTAAGCTGTA<br>AGGATCCTGAGACC |
| G15 | dCas9:VP64 | GGTCTCATATGCCCAAGAAGAAGAGGAAGGTGGAC<br>AAGAAGTACTCCATTGGGCTCGCTATCGGCACAAAC<br>AGCGTCGGCTGGGCCGTCATTACGGACGAGTACAA<br>GGTGCCGAGCAAAAAATTCAAAGTTCTGGGCAATAC<br>CGATCGCCACAGCATAAAGAAGAACCTCATTGGCGC<br>CCTCCTGTTCGACTCCGGGGAAACGGCCGAAGCCA<br>CGCGGCTCAAAAGAACAGCACGGCGCAGATATACC<br>CGCAGAAAGAATCGGATCTGCTACCTGCAGGAGATC<br>TTTAGTAATGAGATGGCTAAGGTGGATGACTCTTTCT<br>TCCATAGGCTGGAGGAGTCCTTTTTGGTGGAGGAG<br>GATAAAAAGCACGAGCGCCACCCAATCTTTGGCAAT<br>ATCGTGGACGAGGTGGCGTACCATGAAAAGTACCC<br>AACCATATATCATCTGAGGAAGAAGCTTGTAGACAG<br>TACTGATAAGGCTGACTTGCGGTTGATCTATCTCGC<br>GCTGGCGCATATGATCAAATTTCGGGGACACTTCCT<br>CATCGAGGGGGACCTGAACCCAGACAACAGCGATG<br>TCGACAAACTCTTTATCCAACTGGTTCAGACTTACAA<br>TCAGCTTTTCGAAGAGAACCCGATCAACGCATCCGG<br>AGTTGACGCCAAAGCAATCCTGAGCGCTAGGCTGTC<br>CAAATCCCGGCGGCTCGAAAACCTCATCGCACAGCT<br>CCCTGGGGAGAAGAAGAACGGCCTGTTTGGTAATCT<br>TATCGCCCTGTCACTCGGGCTGACCCCCAACTTTAA<br>ATCTAACTTCGACCTGGCCGAAGATGCCAAGCTTCA<br>ACTGAGCAAAGACACCTACGATGATGATCTCGACAA |

| | | |
|---|---|---|
| | | TCTGCTGGCCCAGATCGGCGACCAGTACGCAGACC<br>TTTTTTTGGCGGCAAAGAACCTGTCAGACGCCATTC<br>TGCTGAGTGATATTCTGCGAGTGAACACGGAGATCA<br>CCAAAGCTCCGCTGAGCGCTAGTATGATCAAGCGCT<br>ATGATGAGCACCACCAAGACTTGACTTTGCTGAAGG<br>CCCTTGTCAGACAGCAACTGCCTGAGAAGTACAAGG<br>AAATTTTCTTCGATCAGTCTAAAAATGGCTACGCCGG<br>ATACATTGACGGCGGAGCAAGCCAGGAGGAATTTTA<br>CAAATTTATTAAGCCCATCTTGGAAAAAATGGACGG<br>CACCGAGGAGCTGCTGGTAAAGCTTAACAGAGAAG<br>ATCTGTTGCGCAAACAGCGCACTTTCGACAATGGAA<br>GCATCCCCCACCAGATTCACCTGGGCGAACTGCAC<br>GCTATCCTCAGGCGGCAAGAGGATTTCTACCCCTTT<br>TTGAAAGATAACAGGGAAAAGATTGAGAAAATCCTC<br>ACATTTCGGATACCCTACTATGTAGGCCCCCTCGCC<br>CGGGGAAATTCCAGATTCGCGTGGATGACTCGCAAA<br>TCAGAAGAGACTATCACTCCCTGGAACTTCGAGGAA<br>GTCGTGGATAAGGGGGCCTCTGCCCAGTCCTTCAT<br>CGAAAGGATGACTAACTTTGATAAAAATCTGCCTAAC<br>GAAAAGGTGCTTCCTAAACACTCTCTGCTGTACGAG<br>TACTTCACAGTTTATAACGAGCTCACCAAGGTCAAAT<br>ACGTCACAGAAGGGATGAGAAAGCCAGCATTCCTGT<br>CTGGAGAGCAGAAGAAAGCTATCGTGGACCTCCTCT<br>TCAAGACGAACCGGAAAGTTACCGTGAAACAGCTCA<br>AAGAAGATTATTTCAAAAAGATTGAATGTTTCGACTC<br>TGTTGAAATCAGCGGAGTGGAGGATCGCTTCAACGC<br>ATCCCTGGGAACGTATCACGATCTCCTGAAAATCAT<br>TAAAGACAAGGACTTCCTGGACAATGAGGAGAACGA<br>GGACATTCTTGAGGACATTGTCCTCACCCTTACGTT<br>GTTTGAAGATAGGGAGATGATTGAAGAACGCTTGAA<br>AACTTACGCTCATCTCTTCGACGACAAAGTCATGAAA<br>CAGCTCAAGAGGCGCCGATATACAGGATGGGGGCG<br>GCTGTCAAGAAAACTGATCAATGGGATCCGAGACAA<br>GCAGAGTGGAAAGACAATCCTGGATTTTCTTAAGTC<br>CGATGGATTTGCCAACCGGAACTTCATGCAGTTGAT<br>CCATGATGACTCTCTCACCTTTAAGGAGGACATCCA<br>GAAAGCACAAGTTTCTGGCCAGGGGGACAGTCTCC<br>ACGAGCACATCGCTAATCTTGCAGGTAGCCCAGCTA<br>TCAAAAAGGGAATACTGCAGACCGTTAAGGTCGTGG<br>ATGAACTCGTCAAAGTAATGGGAAGGCATAAGCCCG<br>AGAATATCGTTATCGAGATGGCCCGAGAGAACCAAA<br>CTACCCAGAAGGGACAGAAGAACAGTAGGGAAAGG<br>ATGAAGAGGATTGAAGAGGGTATAAAAGAACTGGGG<br>TCCCAAATCCTTAAGGAACACCCAGTTGAAAACACC<br>CAGCTTCAGAATGAGAAGCTCTACCTGTACTACCTG<br>CAGAACGGCAGGGACATGTACGTGGATCAGGAACT |

| | | |
|---|---|---|
| | | GGACATCAATCGGCTCTCCGACTACGACGTGGATG<br>CCATCGTGCCCCAGTCTTTTCTCAAAGATGATTCTAT<br>TGATAATAAAGTGTTGACAAGATCCGATAAAAATAGA<br>GGGAAGAGTGATAACGTCCCCTCAGAAGAAGTTGTC<br>AAGAAAATGAAAAATTATTGGCGGCAGCTGCTGAAC<br>GCCAAACTGATCACACAACGGAAGTTCGATAATCTG<br>ACTAAGGCTGAACGAGGTGGCCTGTCTGAGTTGGAT<br>AAAGCCGGCTTCATCAAAAGGCAGCTTGTTGAGACA<br>CGCCAGATCACCAAGCACGTGGCCCAAATTCTCGAT<br>TCACGCATGAACACCAAGTACGATGAAAATGACAAA<br>CTGATTCGAGAGGTGAAAGTTATTACTCTGAAGTCTA<br>AGCTGGTTTCAGATTTCAGAAAGGACTTTCAGTTTTA<br>TAAGGTGAGAGAGATCAACAATTACCACCATGCGCA<br>TGATGCCTACCTGAATGCAGTGGTAGGCACTGCACT<br>TATCAAAAAATATCCCAAGCTTGAATCTGAATTTGTT<br>TACGGAGACTATAAAGTGTACGATGTTAGGAAAATG<br>ATCGCAAAGTCTGAGCAGGAAATAGGCAAGGCCAC<br>CGCTAAGTACTTCTTTTACAGCAATATTATGAATTTTT<br>TCAAGACCGAGATTACACTGGCCAATGGAGAGATTC<br>GGAAGCGACCACTTATCGAAACAAACGGAGAAACAG<br>GAGAAATCGTGTGGGACAAGGGTAGGGATTTCGCG<br>ACAGTCCGGAAGGTCCTGTCCATGCCGCAGGTGAA<br>CATCGTTAAAAAGACCGAAGTACAGACCGGAGGCTT<br>CTCCAAGGAAAGTATCCTCCCGAAAAGGAACAGCGA<br>CAAGCTGATCGCACGCAAAAAAGATTGGGACCCCAA<br>GAAATACGGCGGATTCGATTCTCCTACAGTCGCTTA<br>CAGTGTACTGGTTGTGGCCAAAGTGGAGAAAGGGA<br>AGTCTAAAAAACTCAAAAGCGTCAAGGAACTGCTGG<br>GCATCACAATCATGGAGCGATCAAGCTTCGAAAAAA<br>ACCCCATCGACTTTCTCGAGGCGAAAGGATATAAAG<br>AGGTCAAAAAAGACCTCATCATTAAGCTTCCCAAGTA<br>CTCTCTCTTTGAGCTTGAAAACGGCCGGAAACGAAT<br>GCTCGCTAGTGCGGGCGAGCTGCAGAAAGGTAACG<br>AGCTGGCACTGCCCTCTAAATACGTTAATTTCTTGTA<br>TCTGGCCAGCCACTATGAAAAGCTCAAAGGATCTCC<br>CGAAGATAATGAGCAGAAGCAGCTGTTCGTGGAACA<br>ACACAAACACTACCTTGATGAGATCATCGAGCAAAT<br>AAGCGAATTCTCCAAAAGAGTGATCCTCGCCGACGC<br>TAACCTCGATAAGGTGCTTTCTGCTTACAATAAGCAC<br>AGGGATAAGCCCATCAGGGAGCAGGCAGAAAACAT<br>TATCCACTTGTTTACTCTGACCAACTTGGGCGCGCC<br>TGCAGCCTTCAAGTACTTCGACACCACCATAGACAG<br>AAAGCGGTACACCTCTACAAAGGAGGTCCTGGACG<br>CCACACTGATTCATCAGTCAATTACGGGGCTCTATG<br>AAACAAGAATCGACCTCTCTCAGCTCGGTGGAGACA<br>GCAGGGCTGATTCGGACCCAAAAAAGAAGCGTAAG |

| | | |
|---|---|---|
| | | GTCGATCCCAAGAAGAAGAGAAAGGTAGATCCTAAGAAGAAGAGAAAGGTAGACGCATTGGATGACTTCGACTTAGACATGCTTGGGTCAGATGCTTTGGACGACTTTGACCTCGACATGTTGGGTTCTGACGCGCTAGATGACTTCGACTTGGACATGTTGGGCTCCGACGCTTTGGACGACTTTGATCTGGACATGTTATGAGGATCCTGAGACC |

### S1.4 gRNA expression cassettes

| Name | Description | Sequence |
|---|---|---|
| U6-gRNA-1 | U6 promoter driving gRNA-1 expression | GGTCTCAAACGTTCGACGTAAAGCCTGTAGAAGAGGTTTCTAGCGAACGACACGAGTTTGAGCCTCATGAAGCTTCGTTGAACAACGGAAACTCGACTTGCCTTCCGCACAATACATCATTTCTTCTTAGCTTTTTTTCTTCTTCTTCGTTCATACAGTTTTTTTTTGTTTATCAGCTTACATTTTCTTGAACCGTAGCTTTCGTTTTCTTCTTTTTAACTTTCCATTCGGAGTTTTTGTATCTTGTTTCATAGTTTGTCCCAGGATTAGAATGATTAGGCATCGAACCTTCAAGAATTTGATTGAATAAAACATCTTCATTCTTAAGATATGAAGATAATCTTCAAAAGGCCCCTGGGAATCTGAAAGAAGAGAAGCAGGCCCATTTATATGGGAAAGAACAATAGTATTTCTTATATAGGCCCATTTAAGTTGAAAACAATCTTCAAAAGTCCCACATCGCTTAGATAAGAAAACGAAGCTGAGTTTATATACAGCTAGAGTCGAAGTAGTGATTAAACCACAATTTGCACACCCGTTTTAGAGCTAGAAATAGCAAGTTAAAATAAGGCTAGTCCGTTATCAACTTGAAAAAGTGGCACCGAGTCGGTGCTTTTTTTGCAAAATTTTCCAGATCGATTTCTTCTTCCTCTGTTCTTCGGCGTTCAATTTCTGGGTTTTTCTCTTCGTTTTCTGTAACTGAAACCTAAAATTTGACCTAAAAAAAATCTCAAATAATATGATTCAGTGGTTTTGTACTTTTCAGTTAGTTGAGTTTTGCAGTTCCGATGAGATAAACCAATAACTTTGCTTAGATCTAATTCATTCCGTTACACCTCTGATGGAGATGGAAGGTTCTTAATAATGATGCCATTTTTTGGGTAATAATTTTGAATTAGAATCAAGGGTATAAGATTCATAATTAACATCACTTAAGCAAAGTTCGTAATATACGACCACAGGATATAATTTTTGGTACGCTGTGAGACC |

| | | |
|---|---|---|
| U6-gRNA-3 | U6 promoter driving gRNA-3 expression | GGTCTCAAACGTTCGACGTAAAGCCTGTAGAAGAGGTTTCTAGCGAACGACACGAGTTTGAGCCTCATGAAGCTTCGTTGAACAACGGAAACTCGACTTGCCTTCCGCACAATACATCATTTCTTCTTAGCTTTTTTTCTTCTTCTTCGTTCATACAGTTTTTTTTTGTTTATCAGCTTACATTTTCTTGAACCGTAGCTTTCGTTTTCTTCTTTTTAACTTTCCATTCGGAGTTTTTGTATCTTGTTTCATAGTTTGTCCCAGGATTAGAATGATTAGGCATCGAACCTTCAAGAATTTGATTGAATAAAACATCTTCATTCTTAAGATATGAAGATAATCTTCAAAAGGCCCCTGGGAATCTGAAAGAAGAGAAGCAGGCCCATTTATATGGGAAAGAACAATAGTATTTCTTATATAGGCCCATTTAAGTTGAAAACAATCTTCAAAAGTCCCACATCGCTTAGATAAGAAAACGAAGCTGAGTTTATATACAGCTAGAGTCGAAGTAGTGATTTGACTCACAGTCCTATCGAGGTTTTAGAGCTAGAAATAGCAAGTTAAAATAAGGCTAGTCCGTTATCAACTTGAAAAAGTGGCACCGAGTCGGTGCTTTTTTTGCAAAATTTTCCAGATCGATTTCTTCTTCCTCTGTTCTTCGGCGTTCAATTTCTGGGTTTTTCTCTTCGTTTTCTGTAACTGAAACCTAAAATTTGACCTAAAAAAAATCTCAAATAATATGATTCAGTGGTTTTGTACTTTTCAGTTAGTTGAGTTTTGCAGTTCCGATGAGATAAACCAATAACTTTGCTTAGATCTAATTCATTCCGTTACACCTCTGATGGAGATGGAAGGTTCTTAATAATGATGCCATTTTTTGGGTAATAATTTTGAATTAGAATCAAGGGTATAAGATTCATAATTAACATCACTTAAGCAAAGTTCGTAATATACGACCACAGGATATAATTTTTGGTACGCTGTGAGACC |
| U6-gRNA-4 | U6 promoter driving gRNA-4 expression | GGTCTCAAACGTTCGACGTAAAGCCTGTAGAAGAGGTTTCTAGCGAACGACACGAGTTTGAGCCTCATGAAGCTTCGTTGAACAACGGAAACTCGACTTGCCTTCCGCACAATACATCATTTCTTCTTAGCTTTTTTTCTTCTTCTTCGTTCATACAGTTTTTTTTTGTTTATCAGCTTACATTTTCTTGAACCGTAGCTTTCGTTTTCTTCTTTTTAACTTTCCATTCGGAGTTTTTGTATCTTGTTTCATAGTTTGTCCCAGGATTAGAATGATTAGGCATCGAACCTTCAAGAATTTGATTGAATAAAACATCTTCATTCTTAAGATATGAAGATAATCTTCAAAAGGCCCCTGGGAATCTGAAAGAAGAGAAGCAGGCCCATTTATATGGGAAAGAACAATAGTATTTCTTATATAGGCCCATTTAAGTTGAAAACAATCTTCAAAAGTCCCACATCGCTTAGATAAGAAAACGAAGCTGAGTTTATATACAGCTAGAGTCGAAGTAGTGATTATTGCGCACCATTCCACTAGGTTTTAGAGCTAGAAATAGCAAGTTAAAATAAGGCTAGTCCGTTATCAACTTGAAAAAGTGGCACCGAGTCGGTGCTTTTTTTGCAAAATTTTCCAGATCGATTTCTTCTTCCTCTGTTCTTCGGCGTTCAATTTCTGGGTTTTTCTCTTCGTTTTCTGTAAC |

| | | |
|---|---|---|
| | | TGAAACCTAAAATTTGACCTAAAAAAAATCTCAAATAATATGATTCAGTGGTTTTGTACTTTTCAGTTAGTTGAGTTTTGCAGTTCCGATGAGATAAACCAATAACTTTGCTTAGATCTAATTCATTCCGTTACACCTCTGATGGAGATGGAAGGTTCTTAATAATGATGCCATTTTTTGGGTAATAATTTTGAATTAGAATCAAGGGTATAAGATTCATAATTAACATCACTTAAGCAAAGTTCGTAATATACGACCACAGGATATAATTTTTGGTACGCTGTGAGACC |
| HHR-gRNA-1 | gRNA-1 expression flanked by 5’ HHR and 3’ HDV | GGTCTCATATGCGACTACTGATGAGTCCGTGAGGACGAAACGAGTAAGCTCGTCAAACCACAATTTGCACACCCGTTTTAGAGCTAGAAATAGCAAGTTAAAATAAGGCTAGTCCGTTATCAACTTGAAAAAGTGGCACCGAGTCGGTGCTTTTGGCCGGCATGGTCCCAGCCTCCTCGCTGGCGCCGGCTGGGCAACATGCTTCGGCATGGCGAATGGGACGGATCCTGAGACC |
| HHR-gRNA-3 | gRNA-3 expression flanked by 5’ HHR and 3’ HDV | GGTCTCATATGGAGTCACTGATGAGTCCGTGAGGACGAAACGAGTAAGCTCGTCTGACTCACAGTCCTATCGAGGTTTTAGAGCTAGAAATAGCAAGTTAAAATAAGGCTAGTCCGTTATCAACTTGAAAAAGTGGCACCGAGTCGGTGCTTTTGGCCGGCATGGTCCCAGCCTCCTCGCTGGCGCCGGCTGGGCAACATGCTTCGGCATGGCGAATGGGACGGATCCTGAGACC |
| HHR-gRNA-4 | gRNA-4 expression flanked by 5’ HHR and 3’ HDV | GGTCTCATATGCGCAATCTGATGAGTCCGTGAGGACGAAACGAGTAAGCTCGTCATTGCGCACCATTCCACTAGGTTTTAGAGCTAGAAATAGCAAGTTAAAATAAGGCTAGTCCGTTATCAACTTGAAAAAGTGGCACCGAGTCGGTGCTTTTGGCCGGCATGGTCCCAGCCTCCTCGCTGGCGCCGGCTGGGCAACATGCTTCGGCATGGCGAATGGGACGGATCCTGAGACC |

**S1.5 Plant expression vector backbone elements**

| Name | Description | Sequence |
|---|---|---|
| Pnos | Nos promoter driving the plant resistance cassette | AGCGGAGAATTAAGGGAGTCACGTTATGACCCCCGCCGATGACGCGGGACAAGCCGTTTTACGTTTGGAACTGACAGAACCGCAACGTTGAAGGAGCCACTCAGCCGCGGGTTTCTGGAGTTTAATGAGCTAAGCACATACGTCAGAAACCATTATTGCGCGTTCAAAAGTCGCCTAAGGTCACTATCAGCTAGCAAATATTTCTTGTCAAAAATGCTCCACTGACGTTCCATAAATTCCCCTC |

| | | |
|---|---|---|
| | | GGTATCCAATTAGAGTCTCATATTCACTCTCAATCC<br>AAATAATCTGCACCGGATCT |
| KanR (Plant) | Kanamycin coding sequence (plant) | ATGATTGAACAAGATGGATTGCACGCAGGTTCTCC<br>GGCCGCTTGGGTGGAGAGGCTATTCGGCTATGAC<br>TGGGCACAACAGACAATCGGCTGCTCTGATGCCG<br>CCGTGTTCCGGCTGTCAGCGCAGGGGCGCCCGGT<br>TCTTTTTGTCAAGACCGACCTGTCCGGTGCCCTGA<br>ATGAACTCCAGGACGAGGCAGCGCGGCTATCGTG<br>GCTGGCCACGACGGGCGTTCCTTGCGCAGCTGTG<br>CTCGACGTTGTCACTGAAGCGGGAAGGGACTGGC<br>TGCTATTGGGCGAAGTGCCGGGGCAGGATCTCCT<br>GTCATCTCACCTTGCTCCTGCCGAGAAAGTATCCA<br>TCATGGCTGATGCAATGCGGCGGCTGCATACGCTT<br>GATCCGGCTACCTGCCCATTCGACCACCAAGCGA<br>AACATCGCATCGAGCGAGCACGTACTCGGATGGA<br>AGCCGGTCTTGTCGATCAGGATGATCTGGACGAA<br>GAGCATCAGGGGCTCGCGCCAGCCGAACTGTTCG<br>CCAGGCTCAAGGCGCGTATGCCCGACGGCGAGGA<br>TCTCGTCGTGACTCATGGCGATGCCTGCTTGCCGA<br>ATATCATGGTGGAAAATGGCCGCTTTTCTGGATTC<br>ATCGACTGTGGCCGGCTGGGTGTGGCGGACCGCT<br>ATCAGGACATAGCGTTGGCTACCCGTGATATTGCT<br>GAAGAGCTTGGCGGCGAATGGGCTGACCGCTTCC<br>TCGTGCTTTACGGTATCGCCGCTCCCGATTCGCAG<br>CGCATCGCCTTCTATCGCCTTCTTGACGAGTTCTT<br>C |
| Tnos | Nos terminator | CTAGAGTCAAGCAGATCGTTCAAACATTTGGCAAT<br>AAAGTTTCTTAAGATTGAATCCTGTTGCCGGTCTTG<br>CGATGATTATCATATAATTTCTGTTGAATTACGTTAA<br>GCATGTAATAATTAACATGTAATGCATGACGTTATT<br>TATGAGATGGGTTTTTATGATTAGAGTCCCGCAATT<br>ATACATTTAATACGCGATAGAAAACAAAATATAGCG<br>CGCAAACTAGGATAAATTATCGCGCGCGGTGTCAT<br>CTATGTTACTAGATCGA |
| pVS1 replicon | OriV; RepA ; StaA for propagation in Agrobacterium | CGTGCGGCTGCATGAAATCCTGGCCGGTTTGTCT<br>GATGCCAAGCTGGCGGCCTGGCCGGCCAGCTTGG<br>CCGCTGAAGAAACCGAGCGCCGCCGTCTAAAAAG<br>GTGATGTGTATTTGAGTAAAACAGCTTGCGTCATG<br>CGGTCGCTGCGTATATGATGCGATGAGTAAATAAA<br>CAAATACGCAAGGGGAACGCATGAAGGTTATCGCT<br>GTACTTAACCAGAAAGGCGGGTCAGGCAAGACGA<br>CCATCGCAACCCATCTAGCCCGCGCCCTGCAACT |

| | | |
|---|---|---|
| | | CGCCGGGGCCGATGTTCTGTTAGTCGATTCCGATC<br>CCCAGGGCAGTGCCCGCGATTGGGCGGCCGTGC<br>GGGAAGATCAACCGCTAACCGTTGTCGGCATCGA<br>CCGCCCGACGATTGACCGCGACGTGAAGGCCATC<br>GGCCGGCGCGACTTCGTAGTGATCGACGGAGCGC<br>CCCAGGCGGCGGACTTGGCTGTGTCCGCGATCAA<br>GGCAGCCGACTTCGTGCTGATTCCGGTGCAGCCA<br>AGCCCTTACGACATATGGGCCACCGCCGACCTGG<br>TGGAGCTGGTTAAGCAGCGCATTGAGGTCACGGA<br>TGGAAGGCTACAAGCGGCCTTTGTCGTGTCGCGG<br>GCGATCAAAGGCACGCGCATCGGCGGTGAGGTTG<br>CCGAGGCGCTGGCCGGGTACGAGCTGCCCATTCT<br>TGAGTCCCGTATCACGCAGCGCGTGAGCTACCCA<br>GGCACTGCCGCCGCCGGCACAACCGTTCTTGAAT<br>CAGAACCCGAGGGCGACGCTGCCCGCGAGGTCC<br>AGGCGCTGGCCGCTGAAATTAAATCAAAACTCATT<br>TGAGTTAATGAGGTAAAGAGAAAATGAGCAAAAGC<br>ACAAACACGCTAAGTGCCGGCCGTCCGAGCGCAC<br>GCAGCAGCAAGGCTGCAACGTTGGCCAGCCTGGC<br>AGACACGCCAGCCATGAAGCGGGTCAACTTTCAGT<br>TGCCGGCGGAGGATCACACCAAGCTGAAGATGTA<br>CGCGGTACGCCAAGGCAAGACCATTACCGAGCTG<br>CTATCTGAATACATCGCGCAGCTACCAGAGTAAAT<br>GAGCAAATGAATAAATGAGTAGATGAATTTTAGCG<br>GCTAAAGGAGGCGGCATGGAAAATCAAGAACAAC<br>CAGGCACCGACGCCGTGGAATGCCCCATGTGTGG<br>AGGAACGGGCGGTTGGCCAGGCGTAAGCGGCTG<br>GGTTGTCTGCCGGCCCTGCAATGGCACTGGAACC<br>CCCAAGCCCGAGGAATCGGCGTGACGGTCGCAAA<br>CCATCCGGCCCGGTACAAATCGGCGCGGCGCTGG<br>GTGATGACCTGGTGGAGAAGTTGAAGGCCGCGCA<br>GGCCGCCCAGCGGCAACGCATCGAGGCAGAAGC<br>ACGCCCCGGTGAATCGTGGCAAGCGGCCGCTGAT<br>CGAATCCGCAAAGAATCCCGGCAACCGCCGGCAG<br>CCGGTGCGCCGTCGATTAGGAAGCCGCCCAAGGG<br>CGACGAGCAACCAGATTTTTTCGTTCCGATGCTCT<br>ATGACGTGGGCACCCGCGATAGTCGCAGCATCAT<br>GGACGTGGCCGTTTTCCGTCTGTCGAAGCGTGAC<br>CGACGAGCTGGCGAGGTGATCCGCTACGAGCTTC<br>CAGACGGGCACGTAGAGGTTTCCGCAGGGCCGGC<br>CGGCATGGCCAGTGTGTGGGATTACGACCTGGTA<br>CTGATGGCGGTTTCCCATCTAACCGAATCCATGAA<br>CCGATACCGGGAAGGGAAGGGAGACAAGCCCGG<br>CCGCGTGTTCCGTCCACACGTTGCGGACGTACTC<br>AAGTTCTGCCGGCGAGCCGATGGCGGAAAGCAGA<br>AAGACGACCTGGTAGAAACCTGCATTCGGTTAAAC |

| | | |
|---|---|---|
| | | ACCACGCACGTTGCCATGCAGCGTACGAAGAAGG CCAAGAACGGCCGCCTGGTGACGGTATCCGAGGG TGAAGCCTTGATTAGCCGCTACAAGATCGTAAAGA GCGAAACCGGGCGGCCGGAGTACATCGAGATCGA GCTAGCTGATTGGATGTACCGCGAGATCACAGAAG GCAAGAACCCGGACGTGCTGACGGTTCACCCCGA TTACTTTTTGATCGATCCCGGCATCGGCCGTTTTCT CTACCGCCTGGCACGCCGCGCCGCAGGCAAGGC AGAAGCCAGATGGTTGTTCAAGACGATCTACGAAC GCAGTGGCAGCGCCGGAGAGTTCAAGAAGTTCTG TTTCACCGTGCGCAAGCTGATCGGGTCAAATGACC TGCCGGAGTACGATTTGAAGGAGGAGGCGGGGCA GGCTGGCCCGATCCTAGTCATGCGCTACCGCAAC CTGATCGAGGGCGAAGCATCCGCCGGTTCCTAAT GTACGGAGCAGATGCTAGGGCAAATTGCCCTAGC AGGGGAAAAAGGTCGAAAAAGCTTCTTTCCTGTGG ATAGCACGTACATTGGGAACCCAAAGCCGTACATT GGGAACCGGAACCCGTACATTGGGAACCCAAAGC CGTACATTGGGAACCGGTCACACATGTAAGTGACT GATATAAAAGAGAAAAAAGGCGATTTTTCCGCCTA AAACTCTTTAAAACTTATTAAAACTCTTAAAACCCG CCTGGCCTGTGCATAACTGTCTGGCCAGCGCACA GCCGAAGAGCTGCAAAAAGCGCCTACCCTTCGGT CGCTGCGCTCCCTACGCCCCGCCGCTTCGCGTCG GCCTATCGCGGCCGCTGGCCGCTCAAAAATGGCT GGCCTACGGCCAGGCAATCTACCAGGGCGCGGAC AAGCCGCGCCGTCGCCACTCGACCGCCGGCGCC CACATCAAGGCACC |
| pMB1 origin | Origin of replication for propagation in E.coli | AAAGGATCTTCCTGAGATCCTTTTTTTCTGCGCGTA ATCTGCTGCTTGCAAACAAAAAAACCACCGCTACC AGCGGTGGTTTGTTTGCCGGATCAAGAGCTACCAA CTCTTTTTCCGAAGGTAACTGGCTTCAGCAGAGCG CAGATACCAAATACTGTCCTTCTAGTGTAGCCGTA GTTAGGCCACCACTTCAAGAACTCTGTAGCACCGC CTACATACCTCGCTCTGCTAATCCTGTTACCAGTG GCTGCTGCCAGTGGCGATAAGTCGTGTCTTACCG GGTTGGACTCAAGACGATAGTTACCGGATAAGGC GCAGCGGTCGGGCTGAACGGGGGGTTCGTGCAC ACAGCCCAGCTTGGAGCGAACGACCTACACCGAA CTGAGATACCTACAGCGTGAGCTATGAGAAAGCGC CACGCTTCCCGAAGGGAGAAAGGCGGACAGGTAT CCGGTAAGCGGCAGGGTCGGAACAGGAGAGCGC ACGAGGGAGCTTCCAGGGGGAAACGCCTGGTATC TTTATAGTCCTGTCGGGTTTCGCCACCTCTGACTT GAGCGTCGATTTTTGTGATGCTCGTCAGGGGGGC GGAGCCTATGGAAAAACGCCAGCAACGCG |

| | | |
|---|---|---|
| KanR (bacterial) | Kanamycin expression cassette for bacterial selection | GCCAATTCGTGCGCGGAACCCCTATTTGTTTATTTTTCTAAATACATTCAAATATGTATCCGCTCATGAGACAATAACCCTGATAAATGCTTCAATAATATTGAAAAAGGAAGAGTATGGCTAAAATGAGAATATCACCGGAATTGAAAAAACTGATCGAAAAATACCGCTGCGTAAAAGATACGGAAGGAATGTCTCCTGCTAAGGTATATAAGCTGGTGGGAGAAAATGAAAACCTATATTTAAAAATGACGGACAGCCGGTATAAAGGGACCACCTATGATGTGGAACGGGAAAAGGACATGATGCTATGGCTGGAAGGAAAGCTGCCTGTTCCAAAGGTCCTGCACTTTGAACGGCATGATGGCTGGAGCAATCTGCTCATGAGTGAGGCCGATGGCGTCCTTTGCTCGGAAGAGTATGAAGATGAACAAAGCCCTGAAAAGATTATCGAGCTGTATGCGGAGTGCATCAGGCTCTTTCACTCCATCGACATATCGGATTGTCCCTATACGAATAGCTTAGACAGCCGCTTAGCCGAATTGGATTACTTACTGAATAACGATCTGGCCGATGTGGATTGCGAAAACTGGGAAGAGGACACTCCATTTAAAGATCCGCGCGAGCTGTATGATTTTTTAAAGACGGAAAAGCCCGAAGAGGAACTTGTCTTTTCCCACGGCGACCTGGGAGACAGCAACATCTTTGTGAAAGATGGCAAAGTAAGTGGCTTTATTGATCTTGGGAGAAGCGGCAGGGCGGACAAGTGGTATGACATTGCCTTCTGCGTCCGGTCGATCAGGGAGGATATCGGGGAAGAACAGTATGTCGAGCTATTTTTTGACTTACTGGGGATCAAGCCTGATTGGGAGAAAATAAAATATTATATTTTACTGGATGAATTGTTTTAGCTGTCAGACCAAGTTTACTCATATATACTTTAGATTGATTTAAAACTTCATTTTTAATTTAAAAGGATCTAGGTGAAGATCCTTTTTGATAATC |
| LB | Left border repeat | CTGATGGGCTGCCTGTATCGAGTGGTGATTTTGTGCCGAGCTGCCGGTCGGGGAGCTGTTGGCTGGCTGGTGGCAGGATATATTGTGGTGTAAACAAATTGACGCTTAGACAACTTAATAACACATTGCGGACGTTTTTAATGTACTG |
| RB | Right border repeat | TGGTTGGCACATACAAATGGACGAACGGATAAACCTTTTCACGCCCTTTTAAATATCCGATTATTCTAATAAACGCTCTTTTCTCTTAGGTTTACCCGCCAATATATCCTGTCAAACACTGATAGTTT |
| GFP DO | GFP expression cassette drop-out flanked by connector sequences | ACTGATGAGACGTGGTAGAGCCACAAACAGCCGGTACAAGCAACGATCTCCAGGACCATCTGAATCATGCGCGGATGACACGAACTCACGACGGCGATCACAGACATTAACCCACAGTACAGACACTGCGACAACGTGGCAATTCGTCGCAATACAACGTGAGACCGAAAGTGAAACGTGATTTCATGCGTCATTTTGAACATTTTGTAAATCTTATTTAATAATGTGTGCGGCAATTCACATTTAATTTATGAATGTTTTCTTAACATCGCGGCAACTCA |

| | | |
|---|---|---|
| | | AGAAACGGCAGGTTCGGATCTTAGCTACTAGAGAA<br>AGAGGAGAAATACTAGATGCGTAAAGGCGAAGAG<br>CTGTTCACTGGTGTCGTCCCTATTCTGGTGGAACT<br>GGATGGTGATGTCAACGGTCATAAGTTTTCCGTGC<br>GTGGCGAGGGTGAAGGTGACGCAACTAATGGTAA<br>ACTGACGCTGAAGTTCATCTGTACTACTGGTAAAC<br>TGCCGGTTCCTTGGCCGACTCTGGTAACGACGCT<br>GACTTATGGTGTTCAGTGCTTTGCTCGTTATCCGG<br>ACCATATGAAGCAGCATGACTTCTTCAAGTCCGCC<br>ATGCCGGAAGGCTATGTGCAGGAACGCACGATTT<br>CCTTTAAGGATGACGGCACGTACAAAACGCGTGC<br>GGAAGTGAAATTTGAAGGCGATACCCTGGTAAACC<br>GCATTGAGCTGAAAGGCATTGACTTTAAAGAGGAC<br>GGCAATATCCTGGGCCATAAGCTGGAATACAATTT<br>TAACAGCCACAATGTTTACATCACCGCCGATAAAC<br>AAAAAAATGGCATTAAAGCGAATTTTAAAATTCGCC<br>ATAACGTTGAGGATGGCAGCGTGCAGCTGGCTGA<br>TCACTACCAGCAAAACACTCCAATCGGTGATGGTC<br>CTGTTCTGCTGCCAGACAATCACTATCTGAGCACG<br>CAAAGCGTTCTGTCTAAAGATCCGAACGAGAAACG<br>CGATCATATGGTTCTGCTGGAGTTCGTAACCGCAG<br>CGGGCATCACGCATGGTATGGATGAACTGTACAAA<br>TGACCAGGCATCAAATAAAACGAAAGGCTCAGTCG<br>AAAGACTGGGCCTTTCGTTTTATCTGTTGTTTGTCG<br>GTGAACGCTCTCTACTAGAGTCACACTGGCTCACC<br>TTCGGGTGGGCCTTTCTGCGTTTATAGGTCTCAGC<br>TGGAAATCTGCTCGTCAGTGGTGCTCACACTGACG<br>AATCATGTACAGATCATACCGATGACTGCCTGGCG<br>ACTCACAACTAAGCAAGACAGCCGGAACCAGCGC<br>CGGCGAACACCACTGCATATATGGCATATCACAAC<br>AGTCCACGTCTCAAGCAGTTACAGAGATGTTACGA<br>ACCG |

**Table S2 : List of OCS constructs used**

| Name | Description | Transcriptional Units |
|---|---|---|
| OCS 1-1 (13,694bp) | U6 driven constitutive expression of gRNA-1 leading to constitutive expression of YFP under the control of pATF-1 promoter | TU1 : U6 :gRNA-1 (1013 bp)<br>TU2 : pATF-1: YFP: T35S (1231 bp)<br>TU3E : P35S : dCas9:VP64: T35S (5162 bp) |

| | | |
|---|---|---|
| OCS 1-5 (13,685bp) | 35S driven expression of gRNA-1 (flanked by ribozymes) leading to constitutive expression of YFP under the control of pATF-1 promoter | TU1 : P35S: HHR-gRNA-1-HDV:T35S (1001 bp)<br>TU2 : pATF-1 : YFP: T35S (1231 bp)<br>TU3E : P35S : dCas9:VP64: T35S (5162 bp) |
| OCS 1-9 (13,592bp) | Ethylene inducible expression of gRNA-1 under the control of EBS promoter. YFP under the control of pATF-1 promoter | TU1 : EBS: HHR-gRNA-1-HDV:T35S (917 bp)<br>TU2 : pATF-1 : YFP: T35S (1231 bp)<br>TU3E : P35S : dCas9:VP64: T35S (5162 bp) |
| OCS 4-1 (14,627bp) | U6 driven constitutive expression of gRNA-1 leading to constitutive expression of Luc under the control of pATF-1 promoter | TU1 : U6 :gRNA-1 (1013 bp)<br>TU2 : pATF-1 : Luc: T35S (2150 bp)<br>TU3E : P35S : dCas9:VP64: T35S (5162 bp) |
| OCS 3-5 (16,989bp) | Ratiometric circuit where YFP under the control of pATF-1 inducible by ethylene, while RFP and BFP under the control of Patf-3 are constitutively expressed | TU1 : EBS: HHR-gRNA-1-HDV:T35S (917 bp)<br>TU2 : pATF-1 : YFP: T35S (1231 bp)<br>TU3: pATF-3 : BFP : T35S (1228 bp)<br>TU4: U6 :gRNA3 (1013 bp)<br>TU5: pATF-3 : RFP : T35S (1214 bp)<br>TU6E : P35S : dCas9:VP64: T35S (5162 bp) |

**Table S3: List of Addgene plasmids used in this study**

| Name | Description |
|---|---|
| pICH86966 Addgene #48075 | Used as the backbone of the shuttle vector; Contains the pVS1 and pMB1 replicons, KanR (bacterial) as well as a plant KanR under pNos. |

| | |
|---|---|
| pYTK001<br>Addgene #65108 | Backbone for cloning all the parts listed in Table S1 |
| pYTK002<br>Addgene #65109 | Contains the connector LS; Used for the construction of TU1 |
| pYTK003<br>Addgene #65110 | Contains the connector L1; Used for the construction of TU2 |
| pYTK004<br>Addgene #65111 | Contains the connector L2; Used for the construction of TU3 |
| pYTK005<br>Addgene #65112 | Contains the connector L3; Used for the construction of TU4 |
| pYTK006<br>Addgene #65113 | Contains the connector L4; Used for the construction of TU5 |
| pYTK007<br>Addgene #65114 | Contains the connector L5; Used for the construction of TU6 |
| pYTK067<br>Addgene #65174 | Contains the connector R1; Used for the construction of TU1 |
| pYTK068<br>Addgene #65175 | Contains the connector R2; Used for the construction of TU2 |

| | |
|---|---|
| pYTK069 Addgene #65176 | Contains the connector R3; Used for the construction of TU3 |
| pYTK070 Addgene #65177 | Contains the connector R4; Used for the construction of TU4 |
| pYTK071 Addgene #65178 | Contains the connector R5; Used for the construction of TU5 |
| pYTK072 Addgene #65179 | Contains the connector RE; Used for the construction of TU6 |
| pYTK095 Addgene #65202 | Used as the backbone for the construction of any transcriptional unit. |
